# Unveiling the Hidden Feast: from molecular detection to predation rate – An example on biological control by generalist predators

**DOI:** 10.1101/2024.11.18.624133

**Authors:** Abel Louis Masson, Kevan Rastello, Ambre Sacco-Martret de Preville, Yann Tricault, Sylvain Poggi, Elsa Canard, Marie-Pierre Etienne, Manuel Plantegenest

## Abstract

Very few processes are as decisive as predation in shaping the structure and dynamics of ecological communities. For most predator species, the number of prey items killed by a predator in a day (predation rate) remains impossible to assess because direct observations are scarce or impossible to acquire. To fill this gap, we propose here a Hierarchical Bayesian Model that integrates data on the molecular detection of prey in predators (e.g. PCR results), on individuals captured in the field on the one hand, and on individuals fed in the laboratory, on the other, in a novel mechanistic framework. By explicitly combining the processes of predation and digestion, model fit provides an estimate of the slope and intercept of the digestion curve, and an estimate of the number of prey consumed by a predator in a day. In a case study targeting 25 carabid beetle species and 5 types of prey in agricultural fields (winter wheat), we use our model to estimate predation rates at species and community scales and demonstrate its advantages for studies on biocontrol and beyond.

Code and Data are available on this repository : <u>/r/HBM_PredationCode-C725/</u>

## 1 Introduction

The trophic ecology of predators shapes the network of interactions within predator-prey assemblages, playing a decisive role in ecosystem functioning and the provision of regulating services such as disease mitigation [O’Bryan et al. 2018] or pest control [Symondson et al. 2002, Bianchi et al. 2006, Snyder et al. 2019]. Prey populations are depleted by a diversity of predators at rates that depend on the diet of each predator species and its predation rate on each type of prey [see Supp. Mat. 0 for definitions]. For some handpicked predators, the per capita predation rate on well-known prey can be measured through direct observations in natural conditions (e.g. lions in the Kruger National Park [Mills and Shenk 1992] or birds foraging on intertidal preys [Wootton 1997]). For most species, however, predation can only be assessed indirectly. This is especially true for species that are difficult to observe whether they are nocturnal, too small, living in remote or inaccessible areas such as telluric species, or because their behavior is highly disrupted by observation. Terrestrial arthropod predators, for example, are ubiquitous but small animals involved in complex prey-predator assemblages in which direct observation of predation events is hardly possible. On the contrary, molecular approaches are well suited to small animals and commonly used to elucidate trophic links [Symondson 2002, King et al. 2008, Traugott et al. 2013, Furlong 2015], but although these are two sides of the same coin, they are rarely used to estimate predation rates. We hypothesize that, despite recent developments (Andow and Paula 2023 & 2024), a methodological breakthrough is still needed to quantify predation levels in complex predator-prey assemblages.

Apart from direct observation, two approaches can be used to obtain estimates of predation rates in real assemblages. The first is to parameterize the functional responses of predators to the density of their prey, typically with lab experiments or in field enclosure conditions [Pawar et al. 2012, Naranjo et Hagler 2001]. But it comes up against two problems. Firstly, lab conditions often lack the complexity of natural environments, such as the presence of multiple prey species, environmental variability, or predator avoidance behaviors, which can significantly affect predation dynamics [Griffen 2021]. Secondly, generalization is difficult when trophic interactions are numerous but poorly understood, which is always the case in large assemblages of inconspicuous species. One way of reducing complexity could be to search for general relationships between the functional traits of predators and/or preys and the corresponding consumption rates, for example by predicting the predation rates using an allometric (body-sized) relationship in a dynamic food web model (Curtsdotter et al., 2019). However, this promising approach requires high-resolution data and raises technical and interpretation problems (Stell et al. 2024). Moreover, parameters are estimated by fitting the entire model to prey population dynamics data, with no possibility of comparing predation rates with other forms of measurement.

A second, more straightforward option is to estimate predation rates using molecular detection of prey ingested by their predators. Since the pioneering work of [Dempster 1960], research has focused on correcting the rough estimate given by the proportion of predators that test positive for prey. Initially simply correcting observed proportions for digestion time [Dempster 1960], models gradually became more complex, integrating, for example, the number of prey eaten when tested positive [Rothschild 1966, Nakamura and Nakamura 1977, Greenstone 1979] or replacing digestion time with the half-life of the prey’s molecular signal in the predator as measured in the lab [Naranjo and Hagler 2001, Greenstone et al. 2010, 2014]. From the 1980s onwards, models associated with quantitative molecular methods [Sunderland 1987, Lister et al. 1987 (b), Sopp et al. 1992, Iverson et al. 2004, Andow and Paula 2023] have replaced the proportion of positive tests with the proportion of prey biomass or DNA remaining in the predator at the time of testing. However, the uncertainty of the measurements provided by quantitative methods is still poorly controlled (Lamb et al. 2019), which could lead to significant prediction errors. For cost reasons, the vast majority of prey molecular detection data remains binary-qualitative (e.g. results of DNA-based methods such as PCR, Multiplex PCR and Metabarcoding; [Furlong 2015, Pereira et al. 2023]). Consequently, another sensible approach is to develop models that make better use of qualitative data, such as Hierarchical Bayesian Models (HBMs) [Coblentz et al. 2017].

Like telescopes for astronomers, models can be used as observation tools to reach hidden information. In particular, models that are anchored in a faithful representation of biological systems allow inferring unobserved quantities through their observed effect on available data. Hierarchical Bayesian modeling [Ellison 2004, Fabre et al. 2010], that naturally accommodates latent variables, provides an accurate framework to develop such an observation tool. Indeed, HBMs allow integrating (i) a representation of all the biological processes of interest through the definition of latent variables, (ii) multiple sources of data relating to all or part of the system, and (iii) all expertise on the system, encompassing both knowledge and uncertainty in the form of *a priori* distributions. To our knowledge, HBMs have never been used to estimate predation rates from molecular data. Yet they seem particularly well suited to the general problem of integrating two sources of data (laboratory and field data) and two processes (digestion and predation) to quantify an underlying variable (predation rate) [Ellison 2004]. Furthermore, unlike frequentist approaches in the literature, they provide a simple and direct quantification of the uncertainties associated with the prediction [Cressie et al. 2009].

In this paper, we present a novel method that uses a hierarchical Bayesian modeling framework to provide estimates of in-field predation rates based on two sources of molecular data: DNA-based prey detection tests carried out on predators fed in the laboratory and on predators captured *in natura*. Simultaneous processing of both datasets enable us to model the gradual extinction of the molecular signal in predators resulting from prey digestion and to estimate prey predation rates, taking into account the digestion process. This new method makes it possible to estimate the effects of the identity of each prey and predator and their interaction on the predation rate. Moreover, our model allows to relax the steady state hypothesis which prevailed in the methods proposed until now [Andow and Paula 2023]. To illustrate the potential benefits of our method, we develop an example of its application to pest regulation by ground beetles (*Carabidae*) in wheat fields. The method successfully provided estimates of predation rates for each predator-prey pair in an assemblage of 25 predator species feeding on 5 types of prey. Thanks to the information transfer enabled by the hierarchical structure of the model, predation rate can be obtained even for predator-prey pairs that are very poorly informed by the data, at the cost of greater uncertainties. The ecological interpretation of the estimated values for the model’s parameters and hyperparameters enhanced our understanding of the biological system. To our knowledge, this is the first example of simultaneously quantifying predation rates of an entire predator community for such a large predator-prey assemblage of soil fauna.

## 2 Model description

Our approach is based on a hierarchical Bayesian model (HBM). All codes mentioned in this section are available in the Supplementary Material. The modeling framework combines a process model, representing the consumption of a prey by a predator through a system of latent variables, and an observation model, which links the estimates of the latent variables to observed data. The process model accounts for a predation process (homogeneous Poisson Process) and a digestion process (exponential decay of prey material in the predator’s gut [Greenstone et al. 2014]). We demonstrate that their combination is an inhomogeneous Poisson Process. It exploits information from two data sources: (i) laboratory data obtained during a feeding trial, consisting of feeding and testing predators at various times after feeding events to assess the decrease in prey DNA detection rate in their gut as a function of time [Greenstone et al. 2014], and (ii) prey DNA detection rate in field-caught predators (Figure 1).

**FIGURE 1.**
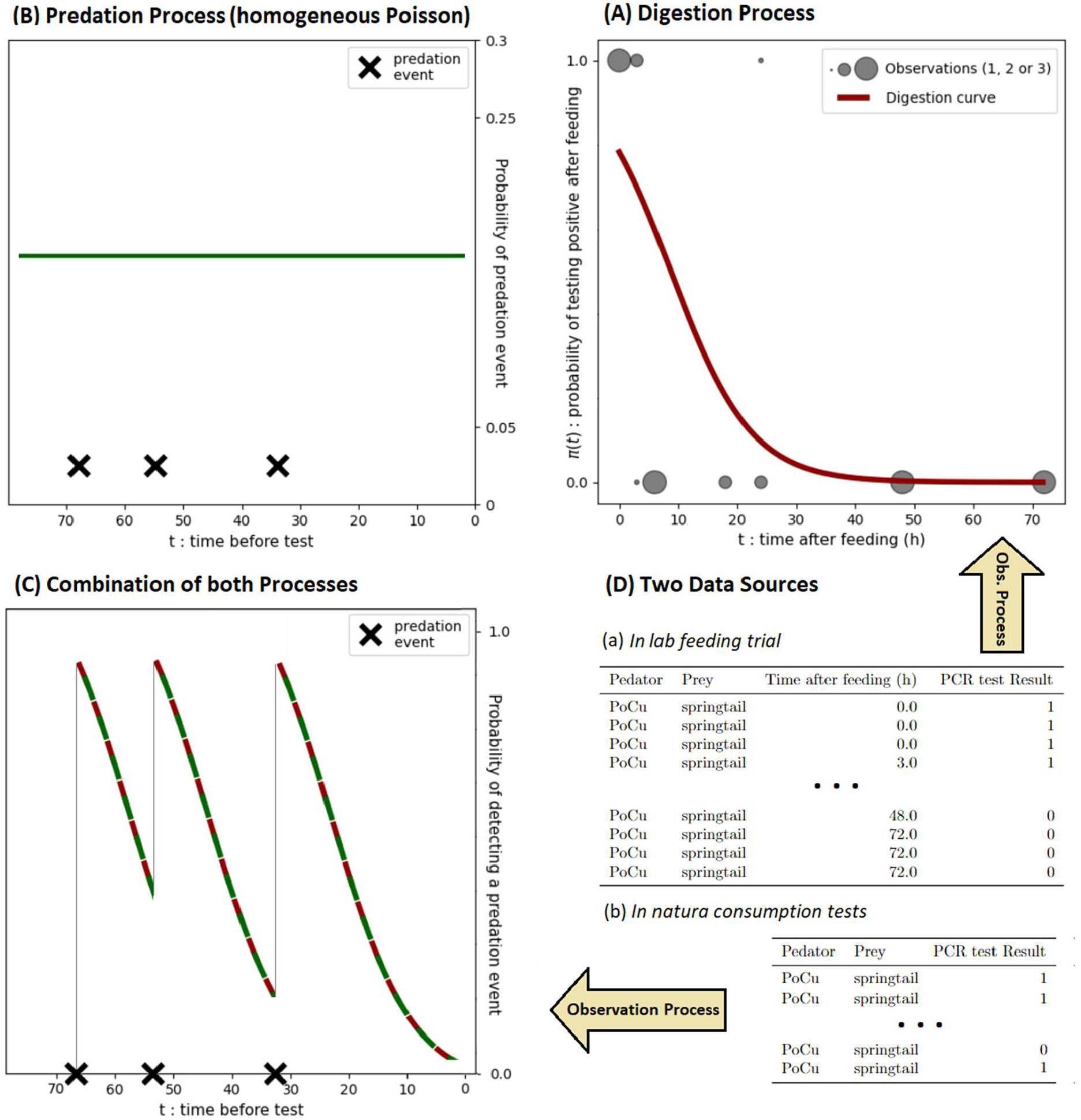
Schematic representation of the model framework. Panel **(A)** shows an example of a digestion curve (in red) compared to the observed data (in gray), which are worth either 1 or zero depending on whether the prey was detected or not (see “PCR test Result” column of Table a in panel (**D)**). Panel **(B)** represents the predation process (in green) with constant predation rate and three predation events occurring in the 72 hours preceding the PCR test. Finally, panel **(C)** represents the combination of predation and digestion (combination of red and green), showing the evolution of the probability of detecting a prey in a predator over time. The data in **(D)** are taken from an example of predation on springtails by the carabid predator Poecilus cupreus (PoCu), as presented in our application example (see 3.1).

### 2.1 Digestion process

Digestion degrades the prey’s DNA, reducing the probability of detection. Let π be the probability of detecting prey’s DNA in the gut content of the predator after a time t since predation. In line with [Greenstone et al. (2014)], we assume that π decreases as a logistic function:

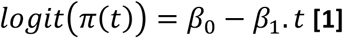

Throughout this document, we refer to the curve defined by π over time as the digestion curve.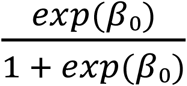 is the probability of detecting prey DNA just after consumption (at the start of the digestion experiment). β_1_ (β_1_ > 0) is the slope of the digestion curve, i.e., the rate at which DNA detection vanishes. It should be noted that the voracity of the predator, i.e. the quantity of prey consumed per meal, is not explicitly accounted for in our model, but depends implicitly on predator, prey and predator-prey interaction (see 2.4).

### 2.2 Predation process

We assume (i) that predation events are independent, (ii) that the predator consumes only one prey item at each predation event, and (iii) that the probability of occurrence of a predation event in a short time interval is proportional to its length. Under these assumptions, the predation process is a homogeneous Poisson process, and *N*_*t*_ the number of predation events during a time interval of length t, follows a Poisson distribution with parameter *λt*, where *λ* is a positive real parameter called the intensity of the Poisson process.

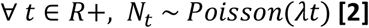

Here, the intensity *λ* is the average number of prey consumed per time step (i.e. per hour), which we refer to as the predation rate.

### 2.3 Observation processes

The first observation process links the expectation provided by the digestion model to the detection tests carried out during the feeding trial. The result of each test 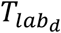 follows a Bernoulli distribution with parameter *π*(*d*), where d is the time elapsed since feeding:

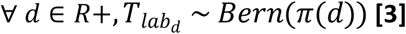

Results of the in-field detection tests also follow a Bernoulli distribution whose parameter is determined by the interactions between predation and digestion processes. A negative detection can occur either from an absence of predation, or because the test failed to detect predation, in particular when digestion has erased the DNA traces.

In [Supp Mat. 2] we demonstrate that *D*, the (theoretical) number of prey that are detected at the time of the test follows a Poisson distribution 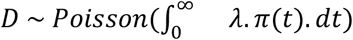.We also show that the integral 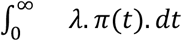 converges and that 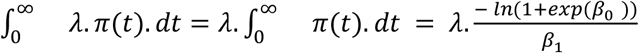

Thus:

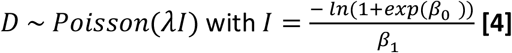

Since we only need one prey to be detected for the test to be positive, the result of a field detection test *T*_*field*_ follows a Bernoulli distribution of probability *pI* = *P*(*D* > 0) = 1 − *P*(*D* = 0).

Furthermore, from equation **[4]** and the Poisson distribution formula we have:

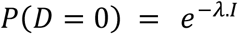

Consequently,

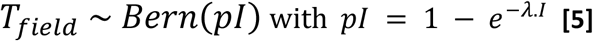

### 2.4 Effects of prey and predator types on predation and digestion processes

#### Parameter decomposition

We assume that the decrease in detection rate with time since feeding depends on the prey, the predator and their combination [Greenstone et al. 2014].

For β_0_, we set:

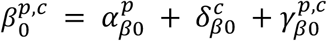

With 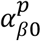,the effect of prey type 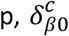,the effect of predator c, and 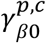,the effect of the prey-predator pair (interaction effect). For the sake of simplicity, we did not distinguish different effects on *β*_1_ and simply used a log-normal distribution to ensure that the detection curve decreases with time since feeding (*β*_1_ > 0 in equation **[1]**) [Supp. Mat. 2].

Similarly, the predation parameters were decomposed into a prey, a predator and a prey-predator interaction effects as follows:

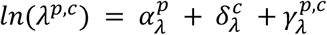

Again the logarithm ensures that the predation rate *λ*^p,c^ remains positive.

#### Hierarchical structure of the model

In the case of strong functional or taxonomic similarities between predator or prey species, some of the prey, predator or predator-prey effects can be hierarchised, allowing, in particular, information to be transferred from well-observed species to those for which observations are scarce. In our application, all predators belong to a single taxonomic group but the number of observations for predator and predator-prey effects varies greatly with many observations for some predator species and very few or even none for others. Hence, parameters 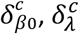 (predator effects), 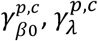 and 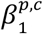 (predator-prey interaction effects) are drawn from normal distributions whose hyperparameters, defined at the level of the entire taxonomic group, indicate the range of their plausible values. On the other hand, as there is no rare prey in the dataset and no taxonomic homogeneity between them, the prey effects are not hierarchised and are assessed independently. The complete structure of the model is shown on Figure 2 below.

**FIGURE 2.**
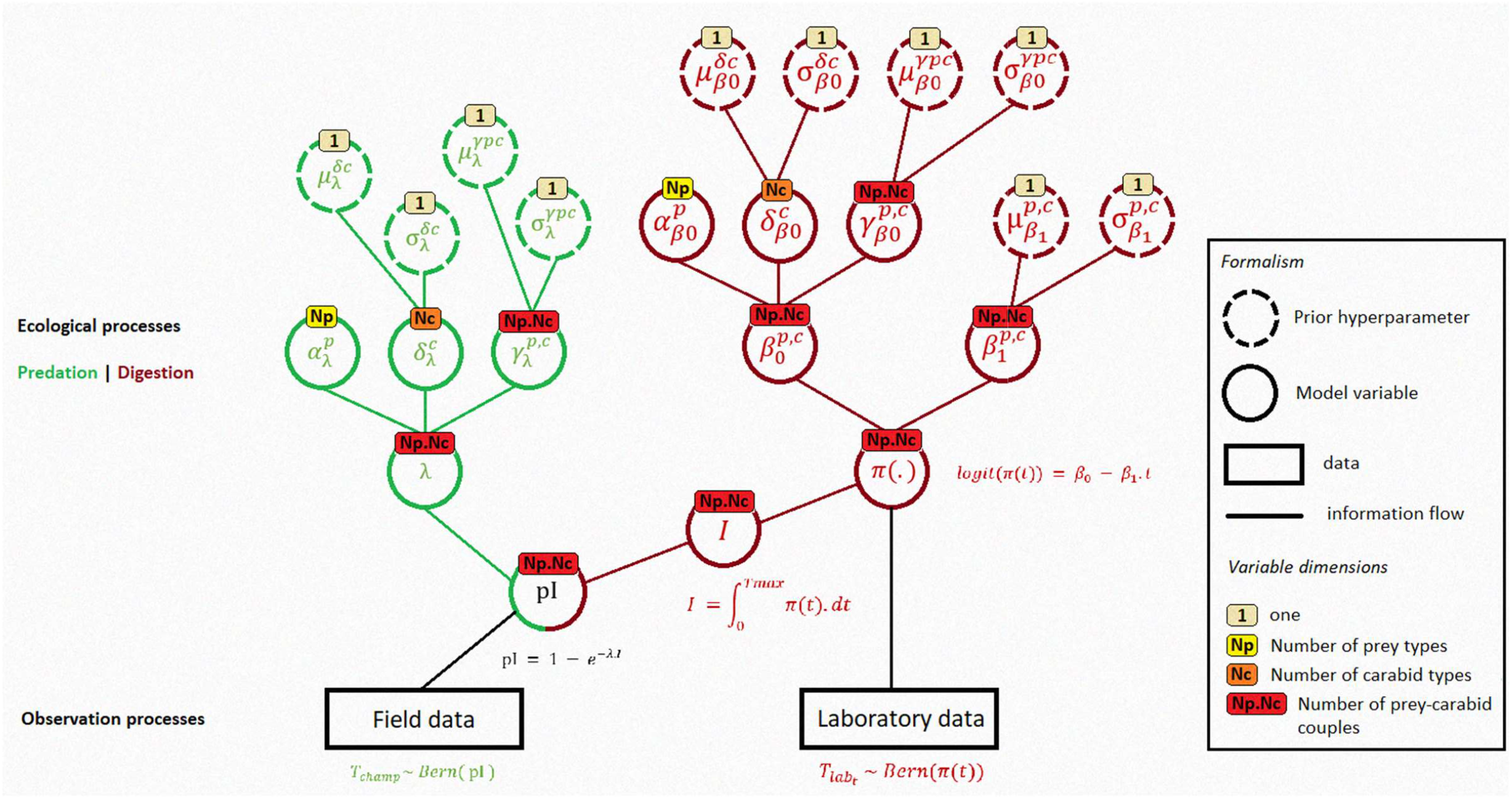
Full model specification. Hyperparameters (dotted circles) are the hyperparameters that we used for our application example (see Supp. Mat. 2. - prior specification). All abbreviations, parameters and variables are defined in Table 1 in Supp. Mat. 1.

## 3 Application example

To illustrate the use of the model, we have applied it to the study of pest regulation by generalist predators, the ground beetles (*Carabidae*), in French wheat fields [Sacco-Martret de Préville et al. 2024]. We focused on the consumption of five types of prey: two groups of pests (slugs and aphids), two groups of decomposers (earthworms and springtails) and one group of natural enemies (spiders, reflecting intraguild predation). As the structure (composition and abundance) of carabid communities is highly seasonal, we used data collected over four sampling sessions [see Supp. Mat. 11]. This example shows how our model can be used to estimate predation rate at the predator species level (see 4.3) and how these results can be integrated at the community level (see 3.2 and 4.4) to provide a new aggregate biocontrol indicator. We also show how our results compare with estimates of predation rates obtained using previously proposed methods (see 4.5).

### 3.1 Data

Two sources of molecular data have been used: one from laboratory feeding experiments and the other from field-caught carabid beetles. Both datasets consist of a set of multiplex PCR results that indicate the presence/absence of the 5 types of prey DNA in the predator’s gut. A total of 542 adults for the feeding trials and of 1534 for the field captures belonging to 25 species of carabid beetles were considered in this study. Field-caught carabids have been collected over 4 sampling sessions in 2018 and 2019 (Autumn (November & December), April, May, and June) and in 4 regions of France (Île-de-France, Bretagne, Pays de la Loire, and Lorraine). For further details on the molecular method used (DNA extraction, primer design, multiplex PCR assay), and on data acquisition please refer to [Sacco-Martret de Préville et al. 2024].

Since [Sacco-Martret de Préville et al. 2024] found that the capture session was the main driver of variation in the structure and activity of carabid communities in the study geographical range, we grouped the individuals caught during the same session in all regions to define four seasonal communities.

### 3.2 Biocontrol indicator at community scale

We introduce a new community-scaled biocontrol indicator as the sum of individual predator contributions to the regulation of each prey:

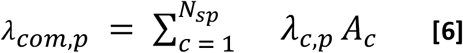

With *N*_*sp*_ the number of carabid species in the community *com, λ*_*c,p*_ the predation rate exerted by carabid species *c* on the considered prey and *A*_*c*_ the abundance (or activity-density) of species *c* in the community.

This biocontrol indicator was calculated for ground beetle communities of Autumn, April, May and June using abundances from each seasonal community but considering constant specific predation rates over the year. For each of the four seasonal community structures, the mean and variance of the indicator were estimated by calculating it with a 100 sets of 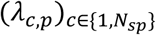 drawn from their joint posterior distributions. For comparison purposes, rough estimates have also been calculated as in [Sacco-Martret de Préville et al. 2024] by replacing the predation rate in equation **[6]** with the proportion of positive detection tests in the observed data.

## 4 Results

### 4.1 Model fitting

Using the priors specified in [Supp. Mat. 2], our model successfully estimates predation parameters for an assemblage of 25 carabid species (i.e. all species present at least once in both feeding experiment and field datasets) and 5 prey types. Convergence of the three Markov chains is validated by Gelman indices values [Gelman et Rubin 1992], all of which are less than 1.01, and visual examination of trace plots (see Figure 8 in Supp. Mat. 3.) and posterior distributions (Figure 14 in Supp. Mat. 6.). Figure 3 illustrates the goodness-of-fit of the model for field detection of springtails [see Figures 9 to 12 in Supp. Mat. 4. for the other prey]. All observed frequencies of detection fall within the 95% interval of the posterior distribution of the detection probabilities (*pI*), and 90% of them stand between the 25th and 75th quantiles. The variability of estimates for each species increases, as expected, as the number of observations decreases, but the estimates remain accurate down to a very small number of observations.

**FIGURE 3.**
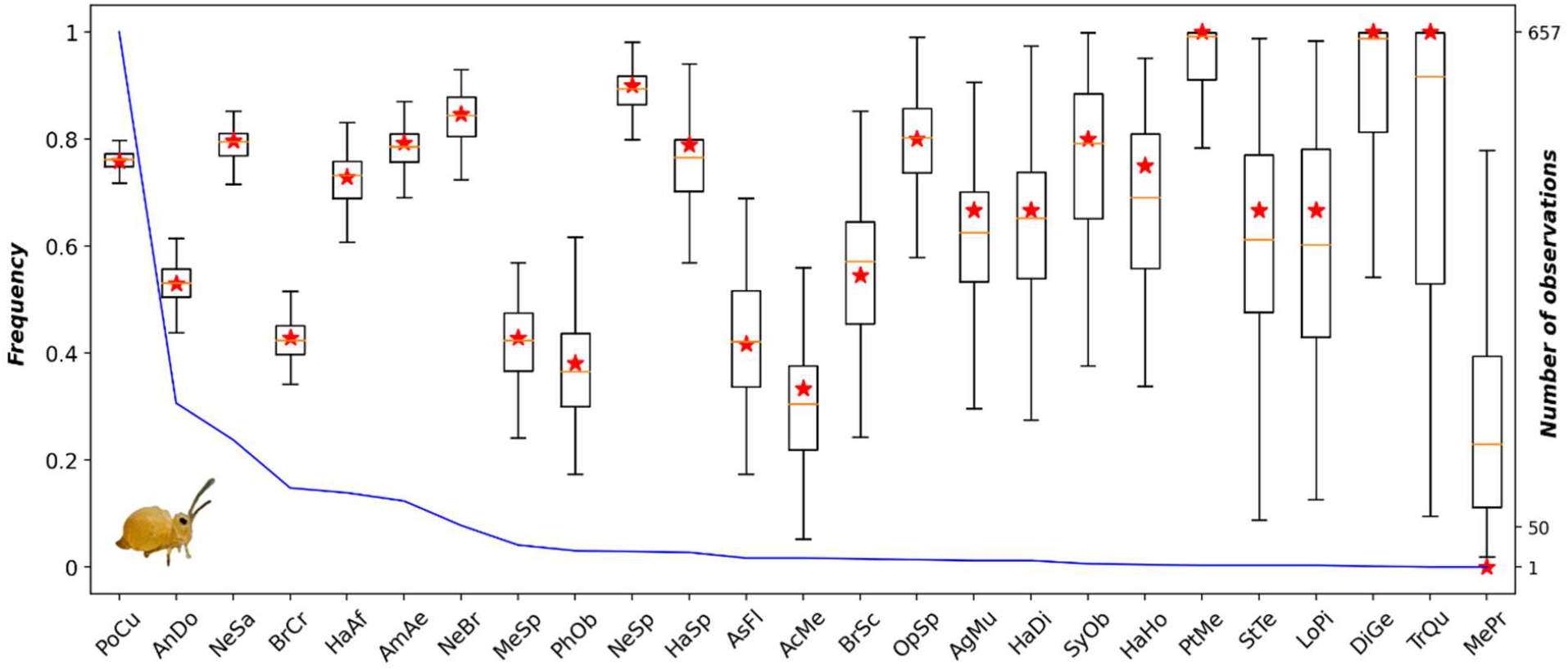
Goodness-of-fit: Posterior distribution of the detection probabilities compared to the observed frequencies of positive tests for each carabid species. Example of springtail detection. Posterior distributions of detection probabilities (pI) are summarized by the boxes, and observed values are indicated by red stars. Carabid species are listed from left to right in descending order of the number of individuals caught in the field (blue curve) ranging from 657 for Poecilus cupreus (PoCu) to 1 observation for Metallina properans (MePr), for a total of 1534 individuals (see Supp. Mat. 10 for carabid code names). The same figures for the other four prey types can be found in Supp. Mat. 4.

### 4.2 Digestion curves

Figure 4 shows that digestion curves vary greatly among the 4 most abundant carabid species in the field and depend on the type of prey consumed. For example, a springtail meal is virtually undetectable 40 hours after feeding in *P. cupreus* **(D)** while more than 72 hours are required for an earthworm meal **(C)**. Similarly, the digestibility of certain prey varies between carabid species. For example, the digestion curves of springtails by *A. aenea, P. cupreus* and *A. dorsalis* are very similar **(D)** while the digestion rate of earthworms or aphids **(C & B)** appear very contrasted between the four species. In particular, for earthworms **(C)**, the digestion curves of *B. sclopeta* and *P. cupreus* differ radically. In *B. sclopeta* the probability of detection is low from the early beginning and decreases rapidly to reach 0 after only 10 hours, whereas in *P. cupreus* the probability of detection is high immediately after feeding (almost 1) and decreases very slowly, still remaining over 0.2 72 hours after ingestion. If we assume that bigger meals take more time to digest, then higher voracity can explain slower digestion rates for *P. Cupreus* and *A. Dorsalis* compared to *B. Sclopeta* (see also Supp. Mat. 5). This example clearly illustrates the need to take digestion curves into account if predation rates are to be correctly estimated from molecular detection data.

**FIGURE 4.**
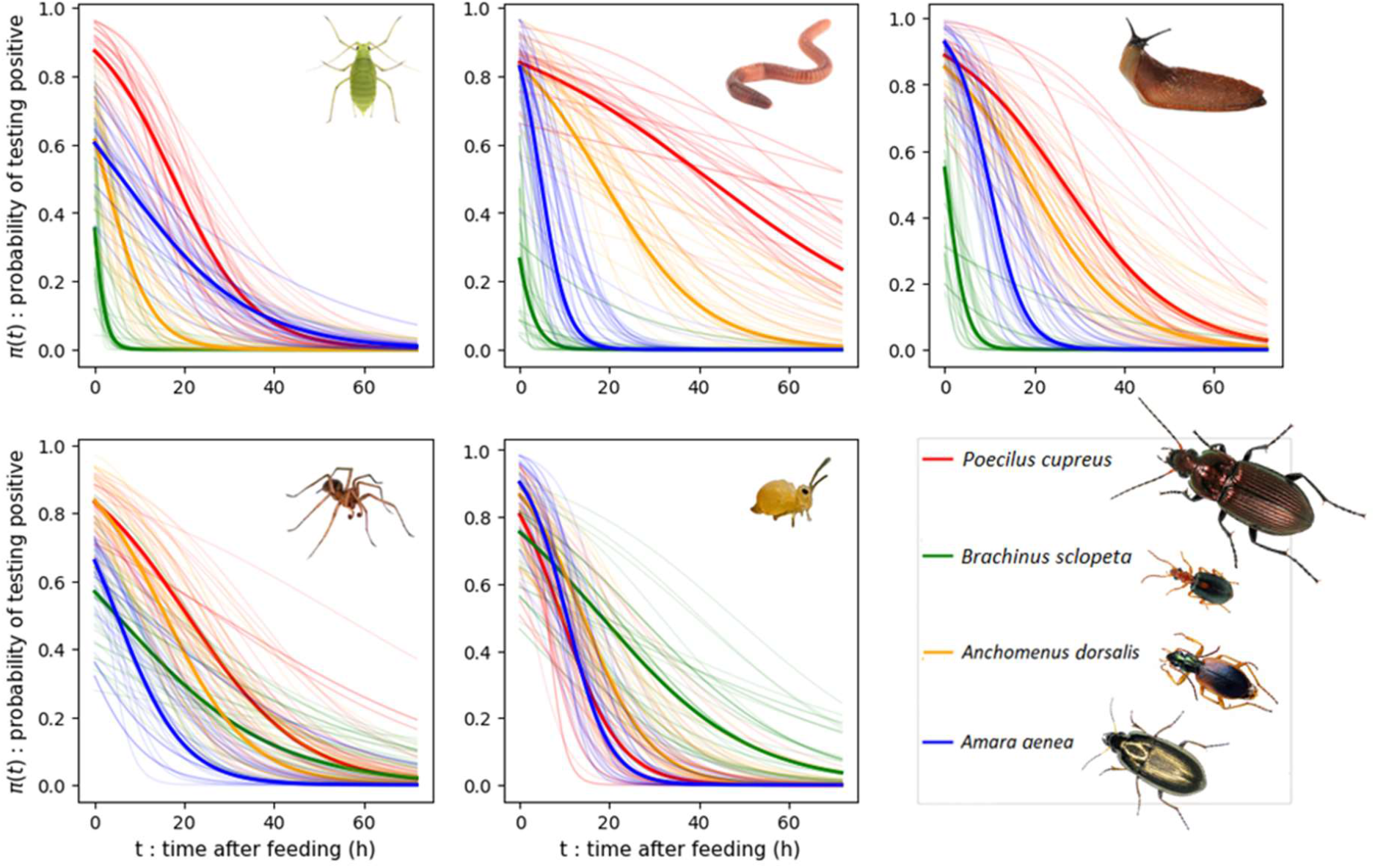
Posterior digestion curves of the five types of prey by the four most abundant carabid species in the dataset. On each panel, the thick bold line is the digestion curve obtained with 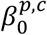 and 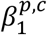 mean posterior values for each predator-prey pair and the 40 shaded lines a random set of posterior curves (i.e. the digestion curves for each predator-prey pair, produced by randomly choosing 40 pairs of 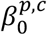 and 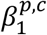 in the joint posterior distribution).

**FIGURE 5.**
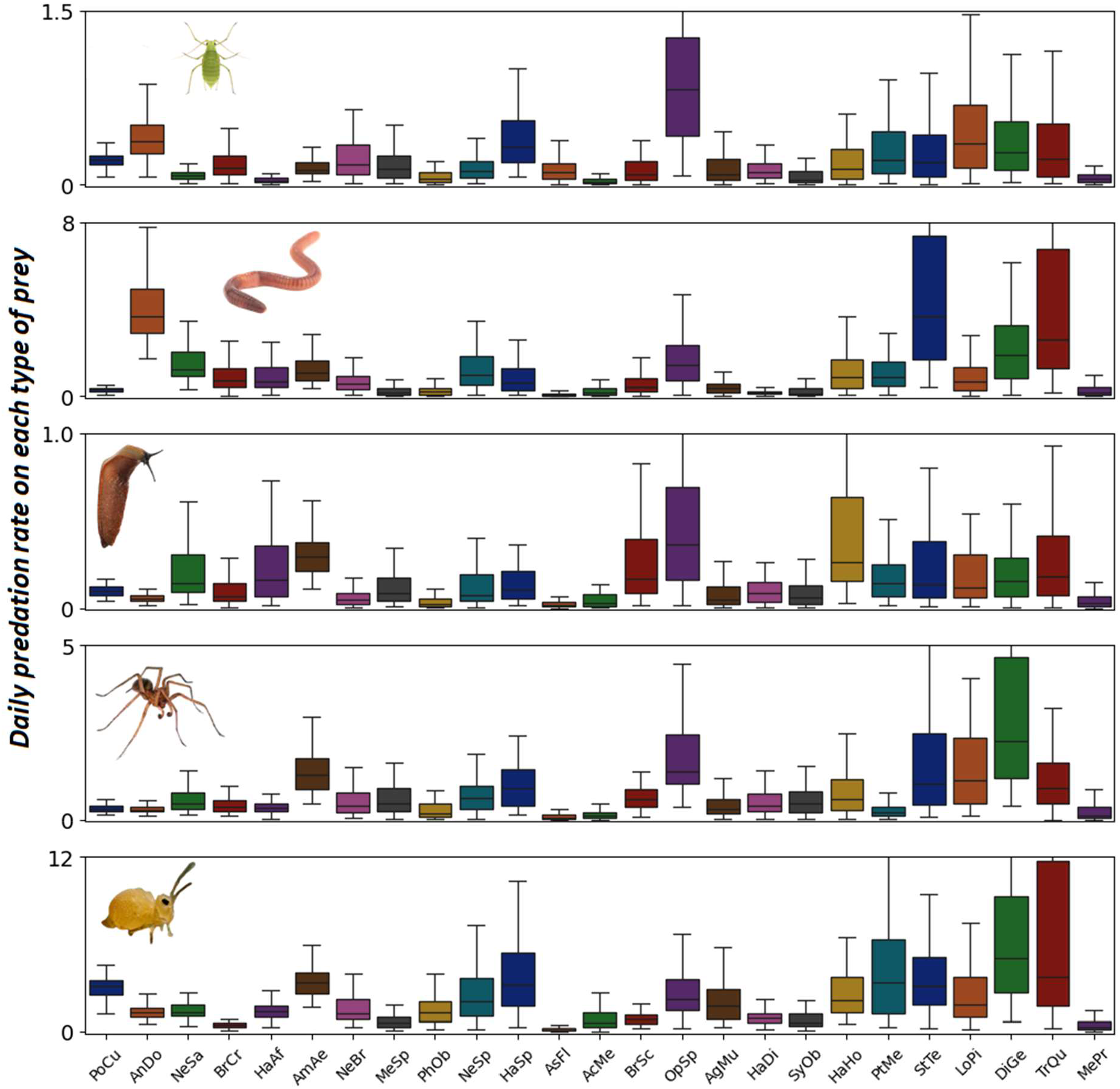
Daily predation rates. Boxplot (median and range) of predation rate estimates on each prey type (i.e. each panel) for each carabid species (x-axis) present in the dataset (see Supp. Mat. 10 for carabid code names). The y-axis range (number of predation events per day) is different for each prey type, as the order of magnitude of predation rate varies between prey types. Colors differ for each carabid species for visual clarity only, and the species are ranked in descending order of abundance in the field. A visualization of the full posterior distributions can be found in Supp. Mat. 6. Posterior distributions of some key prey, predator and interaction effects are also presented in Supp. Mat. 7.

### 4.3 Predation rates

The estimates of predation rate provided by our model fall within a reasonable order of magnitude, both in their median value and in its variability among species. For example, median predation rate on slugs varies from 0.02 feeding event per day for *Phyla obtusa* and *Asaphidion flavipes* to 0.29 for *A. aenea* and 0.36 *for Ophonus sp*. The maximum estimated rate of predation is of the order of 5 prey items consumed per day (median value for predation of springtails by *Diachromus germanus*). For all prey types the median predation rate varies roughly by a factor of ten among predators. This wide range of variation confirms that the biocontrol potential varies greatly according to the composition of the carabid community. Interestingly, distinct predator strategies emerge, with some species favoring predation on a limited number of prey types (e.g. *A. dorsalis* on earthworms and aphids), while many others being true generalists, some even showing high predation rates on all prey types (e.g. *Trechus quadristriatus* and *Stenolophus teutonus*). On the contrary, certain carabid beetles (e.g. *A. flavipes*) appear to be low consumers of the prey studied.

### 4.4 Community-wide biocontrol indicator – Application to the study of seasonal variation

Unsurprisingly, the biocontrol indicator was mainly determined by activity-density, reaching maximum values in May for all prey types. However, the differences in biocontrol indicator values between the autumn and spring communities were less marked for earthworms, slugs and spiders than for aphids and springtails. This result is explained by a change in the composition of the carabid community resulting in a change in food preferences. The seasonal variations of our estimates were similar but more pronounced than those of the rough estimates (red stars on Figure 6) based on species abundances weighted by the detection frequencies (see 3.2). However, the rough estimates systematically underestimated the actual predation rate. The magnitude of the underestimation depended strongly on the prey type considered: it was the lowest for slugs (rough estimates systematically falling between the 25th and 75th quantile of our estimates) and the highest for springtails (rough estimates 3 to 4 times lower than the median of our estimates and systematically below the 99th quantile). This discrepancy can be attributed to variations in digestion curves between prey types. Taking the most abundant *P*.*cupreus* as a reference species, (Figure 4), the difference between the raw prediction and our biocontrol indicator is greater the higher the digestibility of the prey.

**FIGURE 6.**
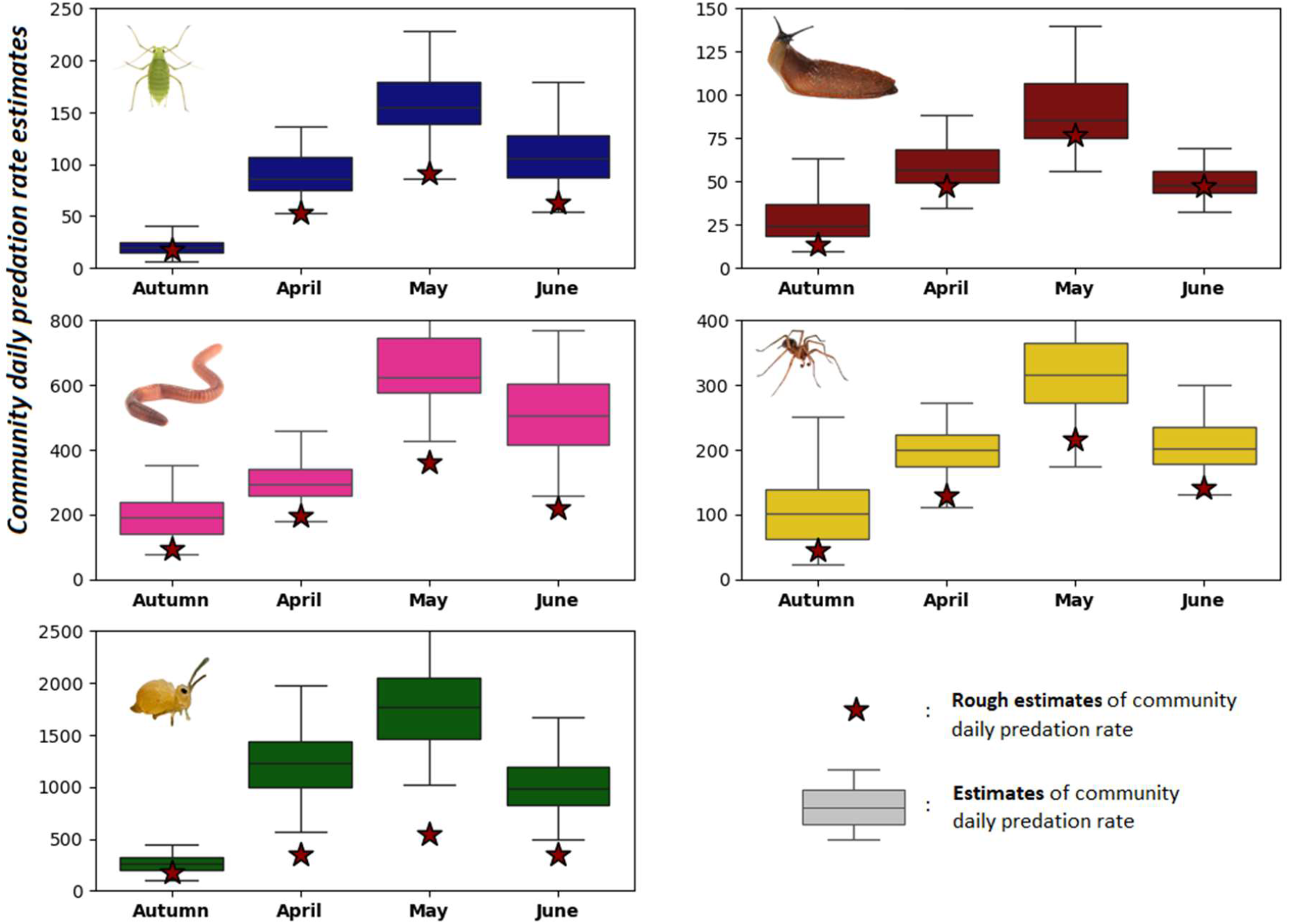
Changes in the biocontrol indicator over the cropping season for the five types of prey. Boxplots show the average predation rate (per day) exerted by the entire community of carabid beetles at each period (x-axis) on each type of prey (panels **(A to E)**). The average predation rate (per day) was calculated according to equation **[6]** (3.2). The red stars are rough estimates, i.e. the sum of species abundances weighted by the detection frequency of each prey (see 3.2).

## 5 Discussion

We have developed a new framework to estimate actual predation rates of inconspicuous predator species in complex terrestrial predator-prey assemblages in the field. It is based on a Hierarchical Bayesian Model mobilizing two sets of binary-qualitative data based on the molecular detection of prey in predators (e.g. PCR results), one in individuals captured in the field, the other in individuals fed in the laboratory.

### 5.1 Performance and biological relevance

The goodness-of-fit of the model to data was very good [Figure 3] but this does not absolutely guarantee the biological relevance of the estimated predation rates. In the absence of direct observations in the field, whose difficulty of access is the very reason for this methodological research, the only values to compare our results with are those obtained by the methods proposed previously or with expert knowledge. The digestion curves obtained in our application example were comparable to the few curves found in the literature [Greenstone et al. 2010, 2014] and in particular to those obtained in a previous study on the same data [Sacco-Martret de Préville et al. 2024]. Our estimates of predation rates were also all within the same orders of magnitude as those obtained with previous methods [see Supp. Mat. 8]. Finally, raw estimates of daily predation rates based solely on detection rates led to values that were systematically lower than those obtained using our model [Figure 6]. Again, this is not surprising given the magnitude of the persistence of the digested prey signal, which in our example is almost always less than one day.

Interestingly, our results were consistent with published information on predator diet. For example, the highest predation rate on earthworms was obtained for a presumed earthworm specialist, *A. dorsalis* [Larochelle 1990]. Similarly, *L. pilicornis*, known in the literature as a springtail specialist [arochelle 1990], showed a high predation rate on this prey. More generally, small carabid beetles generally had higher predation rates than large ones, which is consistent with the fact that they need to feed more frequently. In contrast, the feeding preferences of the carabid beetle *A. aenea*, traditionally described as phytophagous [Kromp 1999], were not confirmed by this study, which focused on animal prey. Its high predation rates on spiders, slugs and springtails combined with good digestion capacities observed in the laboratory rather indicate an omnivorous behavior in our field conditions.

### 5.2 Applicability and conditions of application

The applicability of our model is primarily limited by the ability to collect relevant molecular data on predation and digestion. Limitations arise from the fact that digestion may be affected by the simultaneous consumption of several prey items, and from variations in the predation rate depending on factors such as season or temperature. In addition, molecular methods present inherent limitations, such as the inability to distinguish between dead and live prey, or the stages of prey consumed (eggs, larvae, adults), which have been extensively discussed in the literature [King et al. 2008, Traugott et al. 2013, Furlong 2015]. However, in agreement with the dozen models published on this topic since [Dempster 1960], we believe that these limitations are not substantial enough to prevent the application of the model to a wide range of predators, as well as certain phyto-or coprophagous insects (e.g. granivorous carabid beetles [Carbonne et al. 2020]).

From the 1980s, a major difference in molecular data across studies has been between those indicating prey presence/absence and those offering more detailed information on prey quantity [Sunderland 1987]. The recent model of [Andow and Paula 2023] uses the latter to estimate per capita predation rate from the average quantity of prey DNA in the predators and its rate of decay by digestion, which can be approximated by the inverse of its half-life. The use of quantitative data makes it possible to obtain accurate results with few observations. However the model relies on the assumption that the quantity of prey in predators is constant (steady-state hypothesis, see also [5.4]), which is questionable, especially when the predation rate is low.

In contrast, our model only takes into account the probability of detection, which is better suited to purely qualitative data. Our method is applicable to all binary-qualitative data, provided that **(i)** not all tests are positive, and **(ii)** laboratory and field data have been obtained using the same molecular method and protocol. When the proportion of negative tests is very low, our method reaches saturation and it becomes necessary to measure the amount of prey in the predator, as in [Andow and Paula 2023]. However, as soon as the frequencies of positive and negative tests remain reasonably balanced and the uncertainties and biases associated with the molecular method used are the same for both datasets, our estimates are less sensitive to the molecular method used compared to direct measurements [Andow and Paula 2024].

Finally, the amount of data required to calibrate a model is also crucial to its applicability. Relying on its hierarchy to make the most of each piece of data, we have calibrated our model on a system of 5 prey types x 25 carabid species = 125 interactions, with 1,534 observations in the field and 567 in the laboratory, that is less than 20 observations per predation rate. In [Supp. Mat. 9] we show that it is possible to calibrate the model with even less data.

### 5.3 Potential uses in biocontrol and beyond

As highlighted in [Vialatte et al. 2021], there is an urgent need to quantify pest regulation, especially at the community level of generalist predators that prey on multiple pests. Indeed, there is strong evidence that the abundance and diversity of natural enemy communities promote natural regulation of pests [Vialatte et al. 2021]. However, while the mechanisms involved are generally understood, we still lack the tools to measure them accurately. Some empirical approaches exist, such as the use of exclusion cages [Birkhofer et al. 2017], but these methods are often biased [Furlong et al. 2015] and fail to disentangle the contributions of individual predator species within the community. To the best of our knowledge, the application example on carabid communities represents one of the most comprehensive attempts to quantify the predation rate exerted by an entire predator community on a set of key prey species.

In line with previous work on this issue [Feit et al. 2019, Perennes 2023] we provide a framework that not only inherently considers several prey, but provides predation rate estimates both at species and community scales (see **3.2**). In this way, our model allows not only to estimate the predation rate exerted by the community, which can be translated into a quantification of ecosystem services or disservices depending on the prey consumed (e.g. pest or decomposer), but also to identify, for example, the species that contribute most. Combined with a reliable measure of predator density, which is another challenge [Ahmed et al. 2023], our model provides an unprecedented estimate of the quantity of pests consumed per unit area. Unlike all the proxies and indirect measures currently in use, this estimate would allow direct quantification of the effects of different control strategies (management practices or crop diversification) on pest regulation.

Finally, our model is flexible enough to be used to shed light on more fundamental ecological issues, especially the origins and consequences of predation rate variations. It could be used to analyze the effects of the environment (e.g. season, habitat, prey availability) or predator traits (e.g. size, morphology, diet) on predation rate, as we did with season in [Supp. Mat. 11 and 5.4 below]. One way to do this is to recalibrate the model on subsets of the dataset, as we did in [Supp Mat. 11] for each sampling session. At community scale, our estimates provide valuable information concerning species interactions, such as competition for resources and prey switching which would help us better understand niche differentiation and the determinants of community structure.

### 5.4 Future directions

One simplification of our model, shared with all existing models in the literature, is that predation and digestion rates are constant (steady-state hypothesis). This is obviously not true. However, our model is flexible enough to account for various biotic and abiotic factors that have a proven influence on predation or digestion rates, either by adjusting the hierarchical structure of parameters, or more fundamentally by introducing new variables to overcome the steady-state hypothesis.

A first modification could involve replacing the taxonomic hierarchy used in our study with one based on predator traits (such as size, diet, or reproductive season), which are available for several taxa. A second improvement to the model could focus on parameter decomposition [see 2.4], for instance, by explicitly incorporating the effects of season or prey availability. Indeed, prey availability is strongly influencing predation rates, and in this study, low predation rates on aphids have likely resulted from low aphid densities. Although not directly measured here, previous research has shown mixed effects of prey availability on predation, with some studies finding increased predation at higher prey densities in laboratory conditions [Rothschild 1966, Ge et al. 2019], while others in microcosms, like [Naranjo and Hagler 2001], reported a low impact except at very low densities. In our case, prey density was indirectly considered using the predator’s season of capture as a proxy. In the future, it would be feasible to directly integrate the effect of season or prey density levels into the decomposition of the predation rate [see 2.4].

More substantial modifications to the model can also be considered, particularly to overcome the steady-state hypothesis. [Andow and Paula 2023] proposed a rough linear adjustment of their estimates in cases where predation is not in steady state, based on the increase or decrease of prey biomass divided by the time elapsed between two measurements. We believe that our model is better suited to modifications designed to represent non-stationary situations. Regarding digestion, and since voracity can be measured as part of the feeding experiment [Sacco-Martret de Préville et al. 2024], it is conceivable to integrate it explicitly into the digestion equation. For predation, it is also simple to introduce a variable predation rate, e.g. by making it dependent on temperature or the day/night cycle. These improvements would introduce new sources of variability in detection rates, and each must be tested to ensure they do not prevent the model’s calibration. As an example, in [Supp. Mat. 9], we developed a second version of the model that allows the predation rate to vary depending on the (random) moment when the carabid falls into the pitfall trap, and we show that the model’s calibration remains robust with this modification.

## Supporting information

All Supplementary Materials

Main Code

