## Supplementary material for "Unveiling the Hidden Feast: from molecular detection to predation rate – An example on biological control by generalist predators": All Supplementary Materials

**Supplementary Material 0 |** Glossary

---

**Predator community | Complex of predator | Predator complex**: All predator species coexisting in the same unit of space (e.g. a field) and time (e.g. a cultural season).

**Prey community**: All species preyed upon by the same predator or predator community.

**Predation event**: Killing of prey by predator, with the aim of eating it.

**Predation process**: Accumulation of predation events over time, modeled in our study as a counting process (homogeneous Poisson point process).

**Daily predation rate**: Number of predation events, which we assume corresponds to the number of preys killed over a 24-hour period. In mathematical terms, daily predation rate is the expectation (also known as intensity) of the predation process.

**Digestion (process)**: Degradation of the prey's biomass after a predation event. Represented in our study by the decrease of prey DNA detectability in the predator's gut content.

**Consumption (process)**: Combination of predation and digestion processes.

**Box 1:** Glossary of the key terms used in the description of the modeling framework**.**

**Supplementary Material 1 |** Model parameter and variable description

| | |
|---|---|
| ***Indices*** | |
| $p$ | *Type of prey (taxonomic or functional group)* |
| $c$ | *Type of predator (taxonomic or functional group)* |
| ***Digestion Process*** | |
| $\pi(t)$ | ***Digestion curve**, i.e. probability of prey detection as a function of time after feeding t.* |
| $\beta_0^{p,c}$ | *Y-intercept of the digestion curve* |
| $\beta_1^{p,c}$ | *Slope of the digestion curve* |
| $\alpha_{\beta 0}^{p}$ | *Prey type effect on* $\beta_0^{p,c}$ |
| $\delta_{\beta 0}^{c}$ | *Predator type effect on* $\beta_0^{p,c}$ |
| $\gamma_{\beta 0}^{p,c}$ | *Prey-Predator Interaction effect on* $\beta_0^{p,c}$ |
| $\mu_{\beta_1}^{p,c}$ | *Mean of* $\beta_1^{p,c}$ *distribution* |
| $\sigma_{\beta_1}^{p,c}$ | *Standard deviation of* $\beta_1^{p,c}$ *distribution* |
| ***Predation Process*** | |
| $\lambda^{p,c}$ | ***Predation rate (hourly)**, i.e. amount of prey items of type p predated by predator of type c in an hour.* |
| $\alpha_{\lambda}^{p}$ | *Prey type effect on* $\lambda^{p,c}$ |
| $\delta_{\lambda}^{c}$ | *Predator effect on* $\lambda^{p,c}$ |
| $\gamma_{\lambda}^{p,c}$ | *Prey-Predator Interaction effect on* $\lambda^{p,c}$ |
| ***Observation Processes*** | |
| *Variables* | |
| $N_t$ | *Number of predation events since instant t in the past from which we start counting* |
| $D_t$ | *Number of predation events detected since instant t in the past at the instant of the test* $t = 0$ |
| $T_{lab_t}$ | *Model prediction of PCR test result for a test conducted during the feeding trial at time t* |
| $T_{field}$ | *Model prediction of PCR test result for a test conducted in natura.* |
| *Parameters* | |
| $T_{max}$ | *Time beyond which we consider that predation events can no longer be detected.* |
| $I^{p,c}$ | *Integral of the digestion curve* $\pi$ *over the interval [0, ∞[* |
| $pI^{p,c}$ | *Probability of testing positive to prey of type p for a predator of type c captured in natura.* |

**Table 1***:* Table of parameters & variables

**Supplementary Material 2 |** Mathematical developments

---

***Inhomogeneous Poisson Process***

Let $(D_t)_{t\in R+}$ be the number of predation events occurring during the t hours preceding the test and detected at the time of the test. We assumed that (i) predation event detections are independent and (ii) test success is not influenced by the number of detected predation events. Hence, (iii) the increment in the number of events detected $(D_t)$ between instants $t$ and $t + h$ $(h\ small)$ is equal to the product of the increment in the number of predation events $(N_t)$, which is $\lambda.h$, by the probability that the predated prey DNA can still be detected, which is $\pi(t)$ :

$$\forall\, t > 0,\ D_{t+h} - D_t = \lambda.\pi(t).h + o(h),\ where\ o(h) \rightarrow 0^+ when\ h \rightarrow 0^+$$

Under these three assumptions, the detection process $(D_t)_{t\in R+}$ is an inhomogeneous Poisson process of intensity $\mu(t) = \lambda.\pi(t)$

Thus $D_{t\rightarrow\infty}$ (hereby simply referred to as D) corresponds to the total number of events detected. According to [ref] concerning inhomogeneous Poisson processes, for any $t > 0$, the number of detected predation events $D_t$ follows a Poisson distribution with parameter $\int_0^t \mu(u).du$, so in particular:

$$D \sim Poisson(\int_0^\infty \mu(u).du)\ \textbf{[4]}$$

Finally, a test carried out on a field-caught individual will be positive if at least one predation event is detected, i.e. $D > 0$. Then, the result of a field detection test, $T_{field}$ follows a Bernoulli distribution with a probability $pI = P(D > 0) = 1 - P(D = 0)$. And from equation **[4]** and the Poisson distribution formula (see *Supplementary Material* for a step by step demonstration):

$$P(D = 0) = e^{-\lambda.I} \quad with\ I = \int_0^\infty \pi(t).dt$$

and

$$T_{field} \sim Bern(pI)\ \text{with}\ pI = 1 - e^{-\lambda.I}\ \textbf{[5]}$$

***Prior specification***

For the digestion process, we chose model priors for hyperparameters such that the curves obtained (i) all describe a possible digestion curve, (ii) cover a wide range of digestion curves, and (iii) show no obvious bias towards a particular type of digestion curve (see Figure 7 below).

We chose for each of the 5 prey types a non-informative normal distribution for $\alpha^{p}_{\beta 0}$ and $\alpha^{p}_{\lambda}$:

$\alpha^{p}_{\beta 0} \sim Normal(0, 1)$ and $\alpha^{p}_{\lambda} \sim Normal(0, 20)$

Priors for the hyperparameters of $\delta^{c}_{\beta 0}$, $\delta^{c}_{\lambda}$ (predator effects), $\gamma^{p,c}_{\beta 0}$, $\gamma^{p,c}_{\lambda}$ and $\beta^{p,c}_{1}$ (predator-prey interaction effects) were drawn from normal distributions for the means, and log-uniform distributions for the standard deviations to ensure their strict positivity:

$$\delta^{c}_{\beta 0} \sim Normal(\mu^{\delta c}_{\beta 0}, \sigma^{\delta c}_{\beta 0})\ with\ \mu^{\delta c}_{\beta 0} \sim Normal(0, 0.01)\ and\ ln(\sigma^{\delta c}_{\beta 0}) \sim Uniform(0, 1)$$

$$\gamma^{p,c}_{\beta 0} \sim Normal(\mu^{\gamma pc}_{\beta 0}, \sigma^{\gamma pc}_{\beta 0})\ with\ \mu^{\gamma pc}_{\beta 0} \sim Normal(0, 0.01)\ and\ ln(\sigma^{\gamma pc}_{\beta 0}) \sim Uniform(0, 1)$$

$$ln(\beta^{p,c}_{1}) \sim Normal(\mu^{p,c}_{\beta_1}, \sigma^{p,c}_{\beta_1})\ with\ \mu^{p,c}_{\beta_1} \sim Normal(-2, 0.01)\ and\ ln(\sigma^{p,c}_{\beta_1}) \sim Uniform(0, 1)$$

$$\delta^{c}_{\lambda} \sim Normal(\mu^{\delta c}_{\lambda}, \sigma^{\delta c}_{\lambda})\ with\ \mu^{\delta c}_{\lambda} \sim Normal(0, 0.01)\ and\ ln(\sigma^{\delta c}_{\lambda}) \sim Uniform(0, 100)$$

$$\gamma^{p,c}_{\lambda} \sim Normal(\mu^{\gamma pc}_{\lambda}, \sigma^{\gamma pc}_{\lambda})\ with\ \mu^{\gamma pc}_{\lambda} \sim Normal(0, 0.01)\ and\ ln(\sigma^{\gamma pc}_{\lambda}) \sim Uniform(0, 100)$$

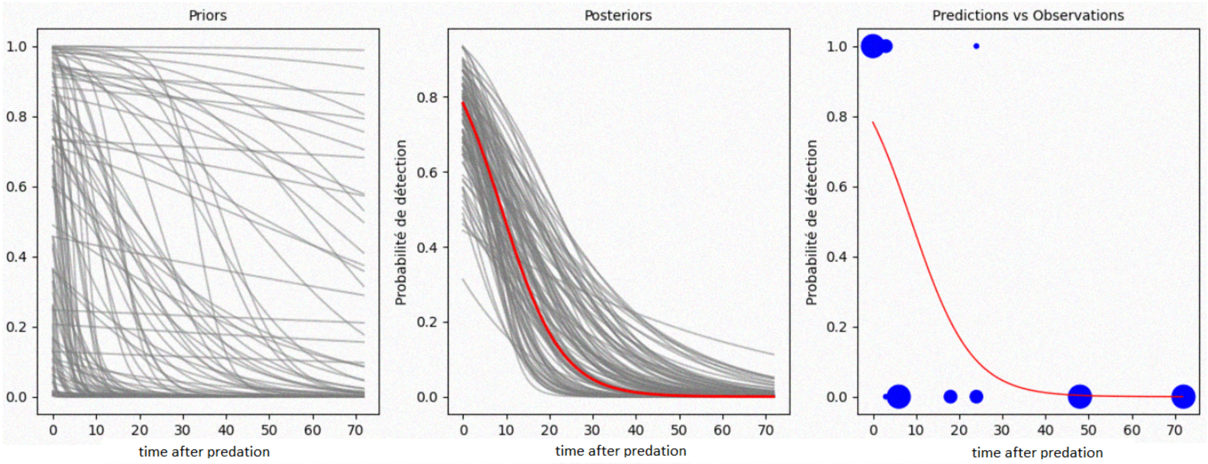


**FIGURE 7** – ***Example of aphid digestion by Poecilus Cupreus***. *The sub-figures show a priori digestion curves (column 1), a posteriori digestion curves (column 2), and the average a posteriori digestion curve compared with the observed data (column 3). The 100 grey curves shown in column 1 (resp. 2) were obtained by drawing parameter values randomly – yet jointly - from their priors (resp. posteriors). The figure on the right compares the average a posteriori digestion curve (in red) with observations (detections in blue) as a function of time. The size of the dots corresponds to the number of observations from 1 to 3.*

***Analytic expression of Integral*** $I$

For any given T > 0, we have :

$$I(T) \; = \; \int_0^T \frac{exp(\beta_0 + \beta_1.t)}{(1+exp(\beta_0 + \beta_1.t))} \, dt$$

We pose, $u \; = \; \beta_0 \; + \beta_1 . t$

$$I(T) = \int_{\beta 0}^{\beta 0 + \beta_1 .T} \frac{1}{\beta_1} \frac{exp(u)}{(1+exp(u))} du$$

$$= \frac{1}{\beta_1} [ln(1 \; + \; exp(u))]_{\beta_0}^{\beta_0 + \beta_1 .T}$$

Hence,

$$I(T) = \frac{ln(1+exp(\beta_0 + \beta_1 .T)) - ln(1+exp(\beta_0))}{\beta_1^{p,c}}$$

Finally, as $I \; = \lim_{T \to \infty} \; I(T)$ and $\lim_{T \to \infty} ln(1 \; + \; exp(\beta_0 \; + \beta_1 . T)) \; = \; 0$ we deduce that $I$ converges and that:

$$I \; = \; \frac{- \, ln(1+exp(\beta_0))}{\beta_1}$$

Graphical analysis of convergence, as well as convergence metrics values are available in the documented code for all samples presented in this study.

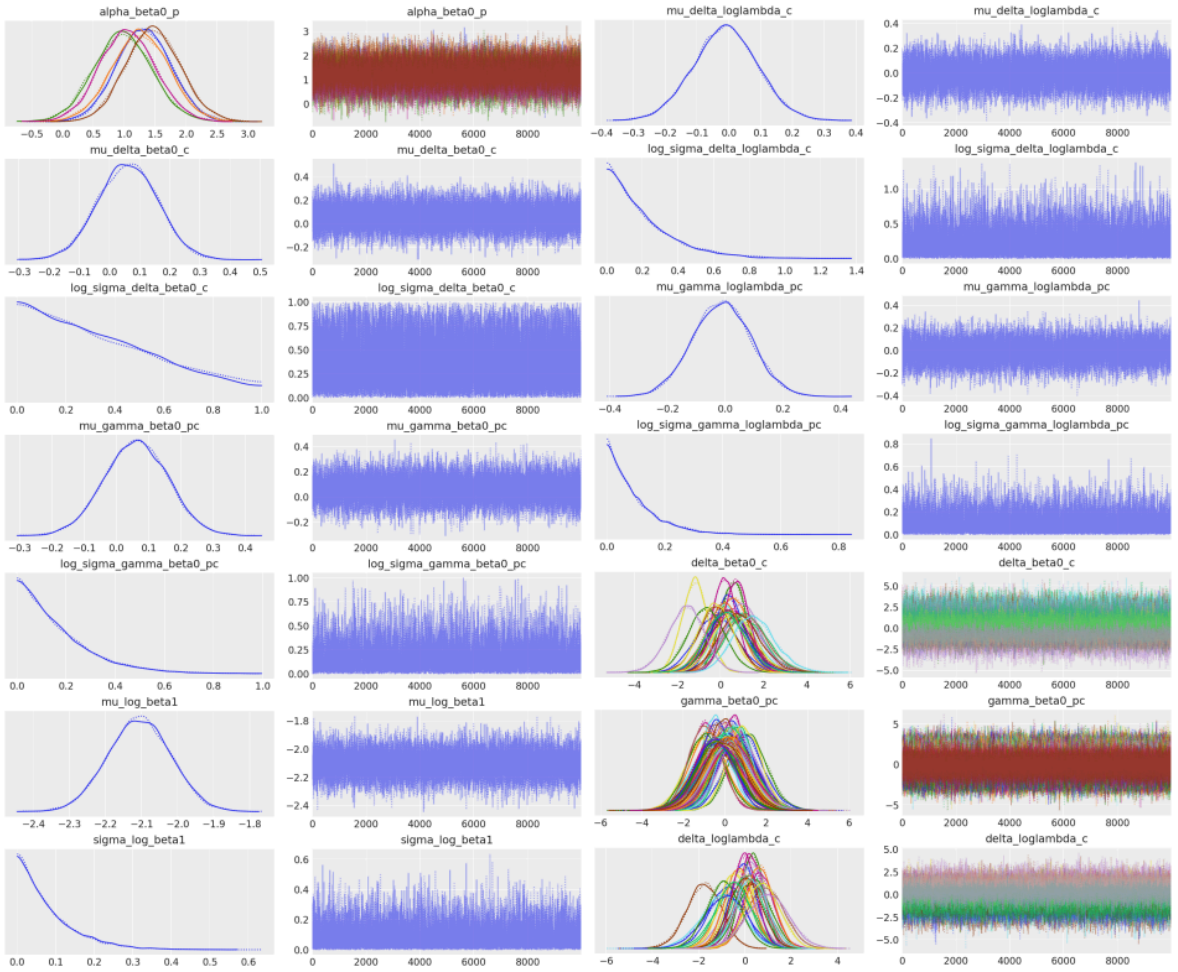


**Figure 8** : Posterior distributions (left panels) and markov chains (right panels) for the main parameters as simulated in the sample *Sample_Predation_Model.nc*

**Supplementary Material 4 |** Model goodness-of-fit– Estimates vs. observations of the frequency of positive tests

All four Figures 9 – 12 share the same legend as Figure 3. The only difference between each of them lies in the prey that is considered.

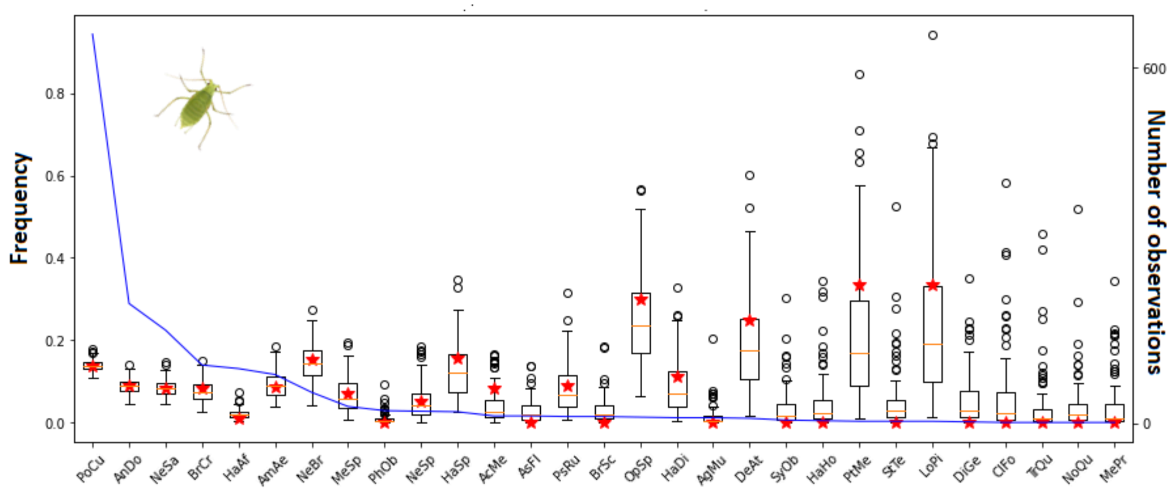


**Figure 9** - Estimates vs. observations of the frequency of positive tests for aphids

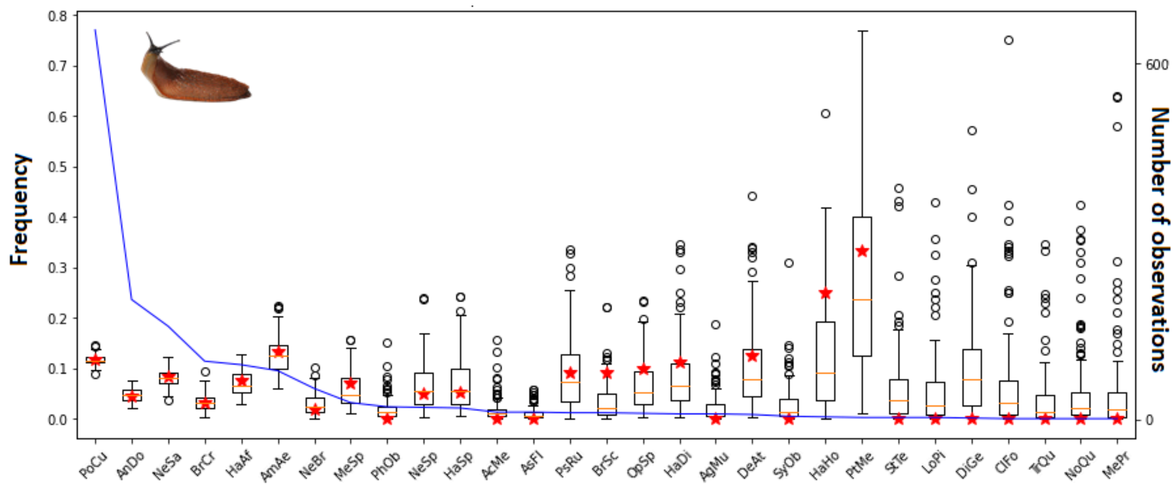


**Figure 10** - Estimates vs. observations of the frequency of positive tests for slugs

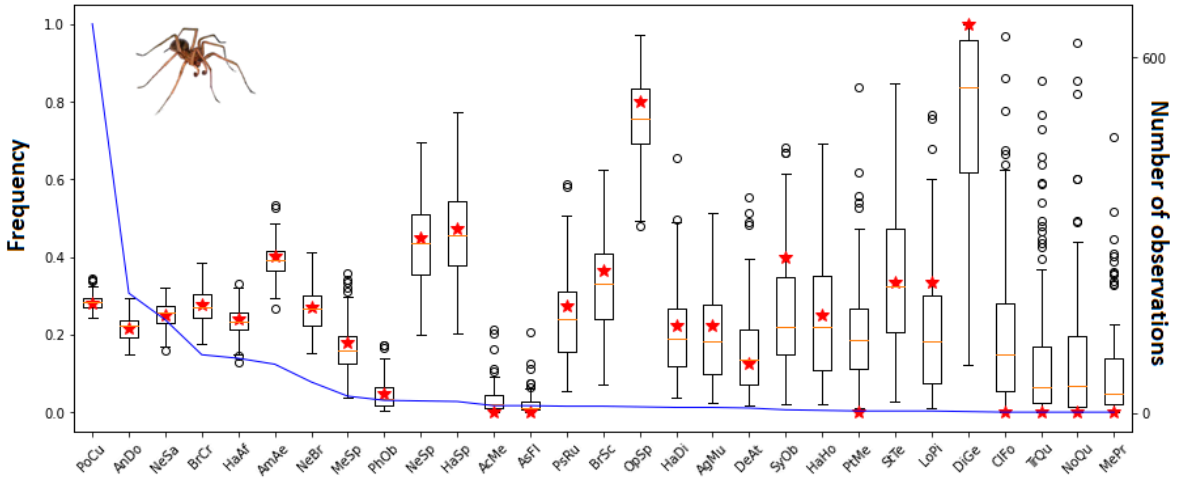


**Figure 11** - Estimates vs. observations of the frequency of positive tests for spiders

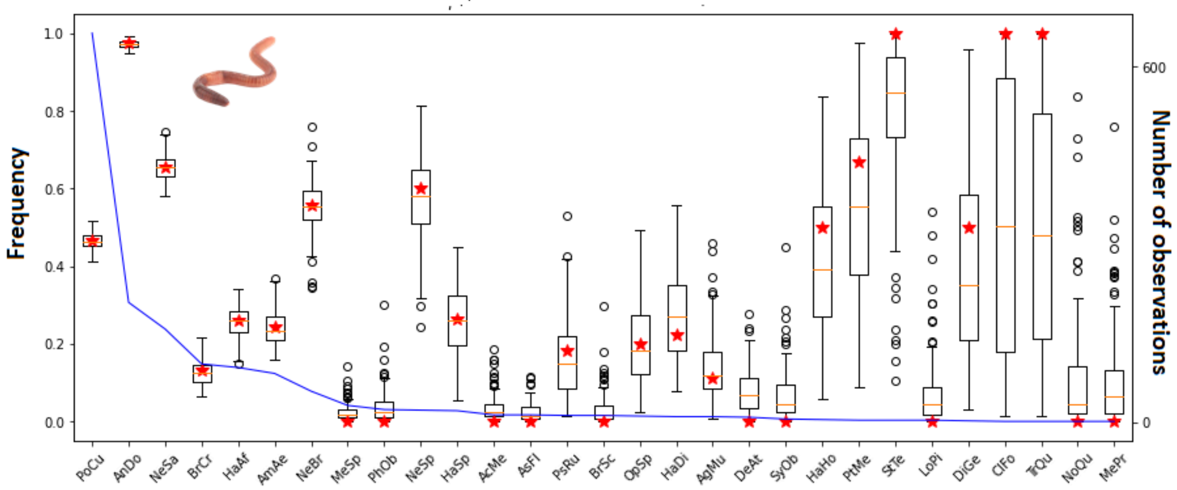


**Figure 12** - Estimates vs. observations of the frequency of positive tests for earthworms

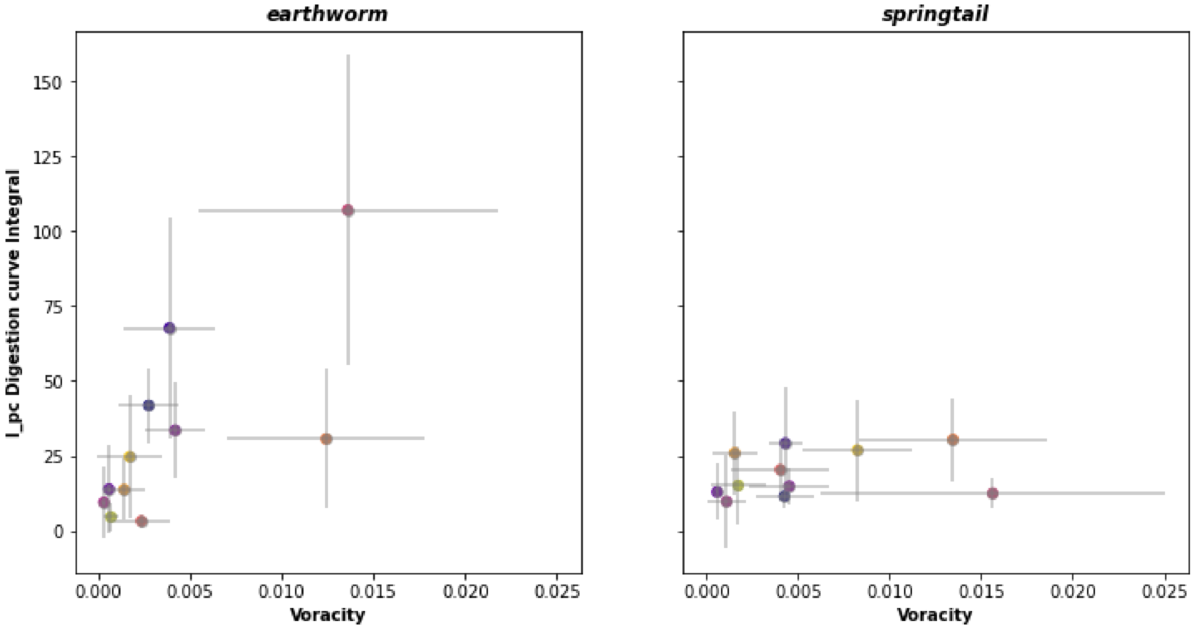


**FIGURE 13** – *Mean voracity (measured as presented in [Sacco Martret de Préville et al. 2024]) against mean digestion properties (predicted). The predictions are given in the form of digestion curve integral values for each carabid species on earthworm (panel* ***A****) and springtails (panel* ***B****).*

**Supplementary Material 6 |** Posterior distributions of daily predation rate for each predator-prey pair

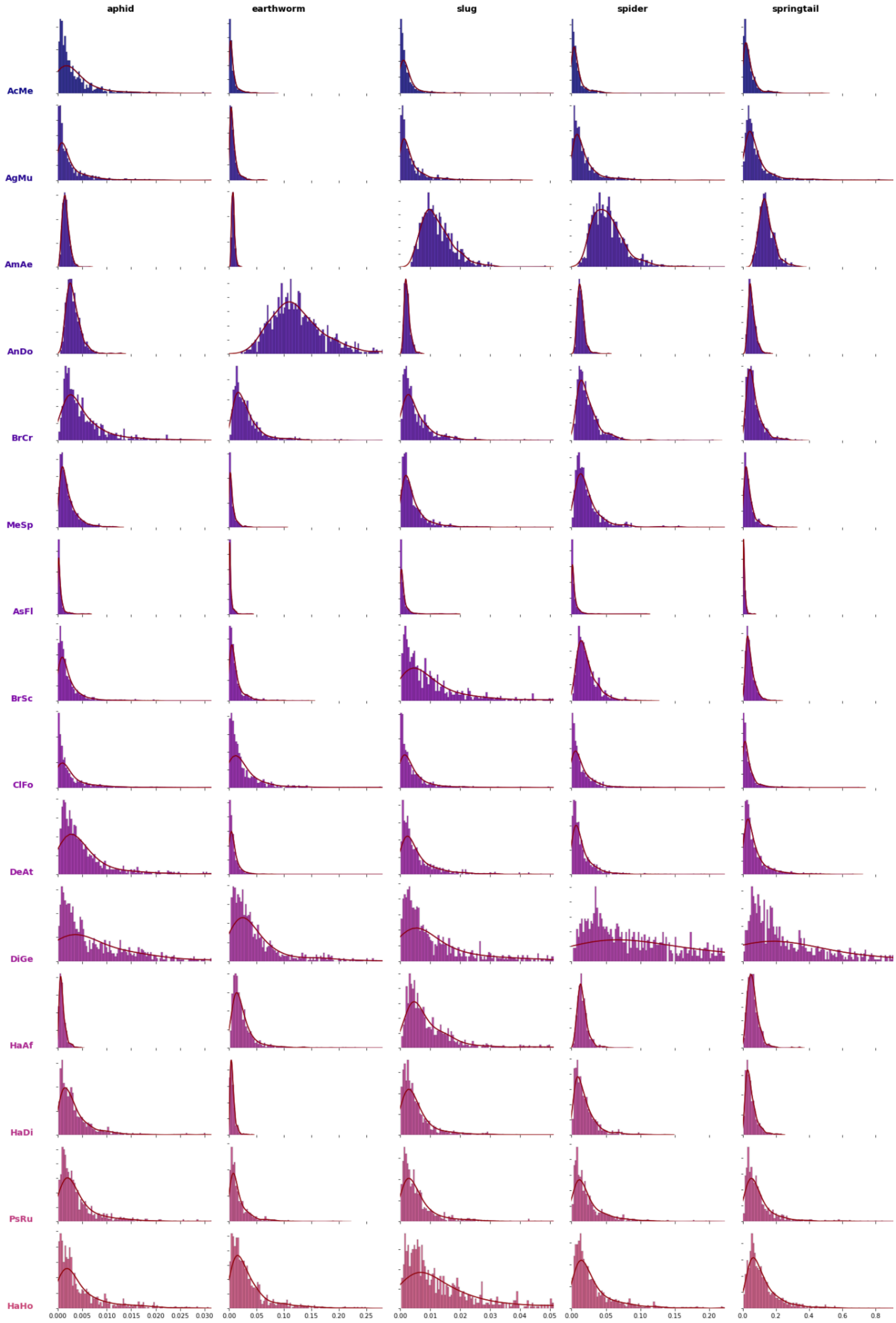

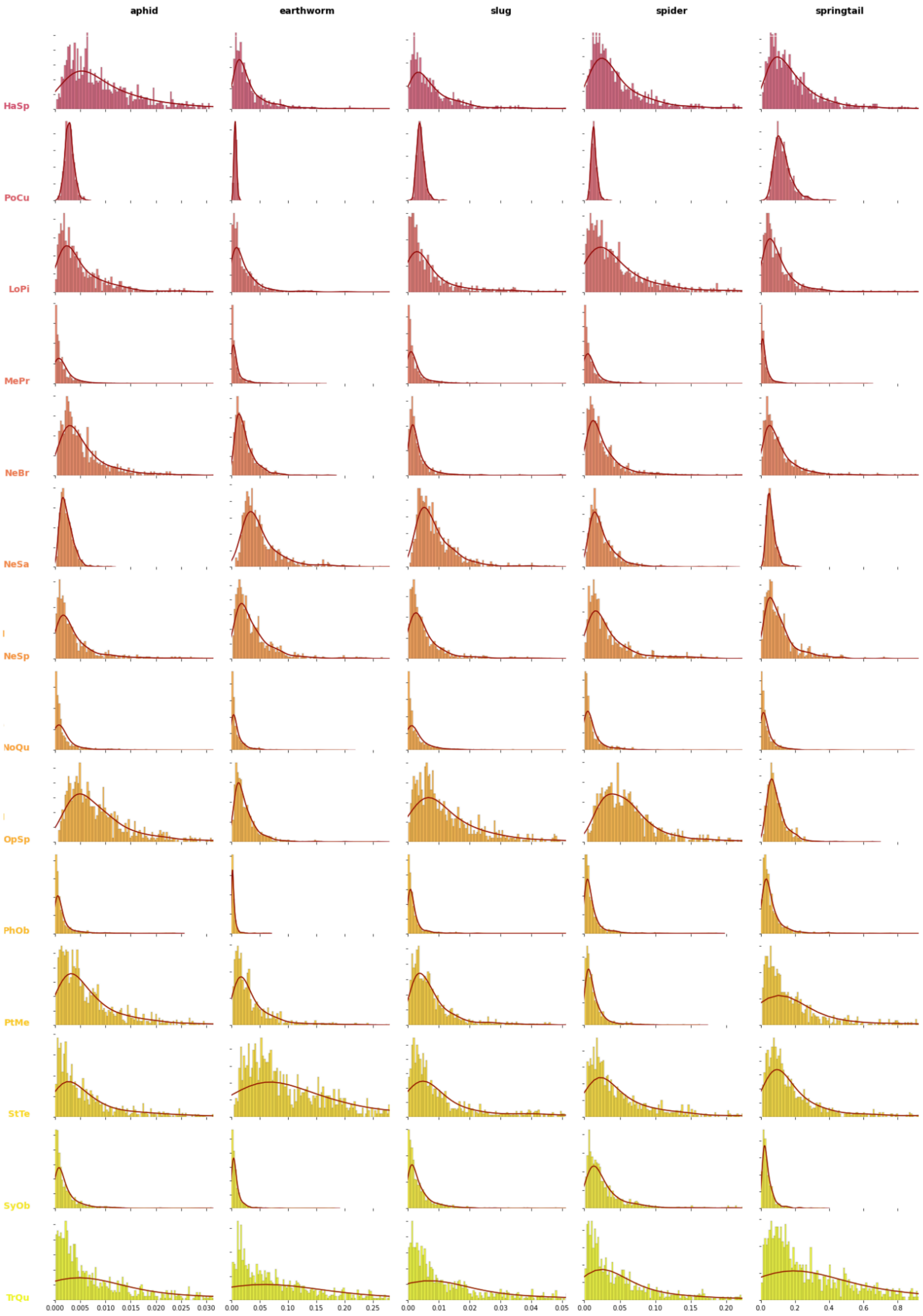


**FIGURE 14** – *Posterior Distributions of predation rate lambda for the 29 carabid species of the dataset on each of the five types of preys. Color coding is for visual clarity only. The abscissa is*

*different for each prey type (i.e. each column), since the order of magnitude of the number of predation events is different for each prey type.*

Despite the convergence of the Markov Chains, the model's estimates naturally reflect the imbalance in the number of observations for each carabid species *in natura* (from 657 observations for *Poecilus Cupreus* to 1 for *Trechus Quadristriatus*) or under controlled conditions (between 66 and 101 for the four main species presented below, to 1 for the least represented species e.g. *Pseudophonus Rufipes*). In particular, in the case of *Trechus Quadristriatus* this imbalance resulted in posteriors that improved on priors, yet remained very flat (see Figure 14).

The posteriors provided by the model have a reasonable shape (no bi- or tri-modality), and a manageable variance, even for species for which little data is available. The variance seems to increase with the average predation intensity, which is best exemplified in the case of *Poecilus Cupreus* showing a much higher mean and variance on springtails compared to the other preys. However, it seems that saturation is never reached, meaning that predation intensity never got so high that it became impossible to estimate. Even the very flat posteriors of *Trechus Quadristriatus* are within reason.

**Supplementary Material 7 |** Prey, Predator and Interaction effects

As mentioned in the introduction, our model is anchored in a realistic representation of predator-prey interactions. As such, most of its parameter's distributions can be interpreted biologically to provide information about the prey, the predator or their interaction. Typically, predation rate $\lambda_{pc}$ is decomposed into a **prey effect** $\alpha_{\lambda}^{p}$, which informs us about the **relative frequency of predation events** for each prey, a **predator effect** $\delta_{\lambda}^{c}$ accounting for the **predator's voracity**, and an **interaction effect** $\gamma_{\lambda}^{p,c}$ that reflects the **degree of specialization** of the predator for the prey.

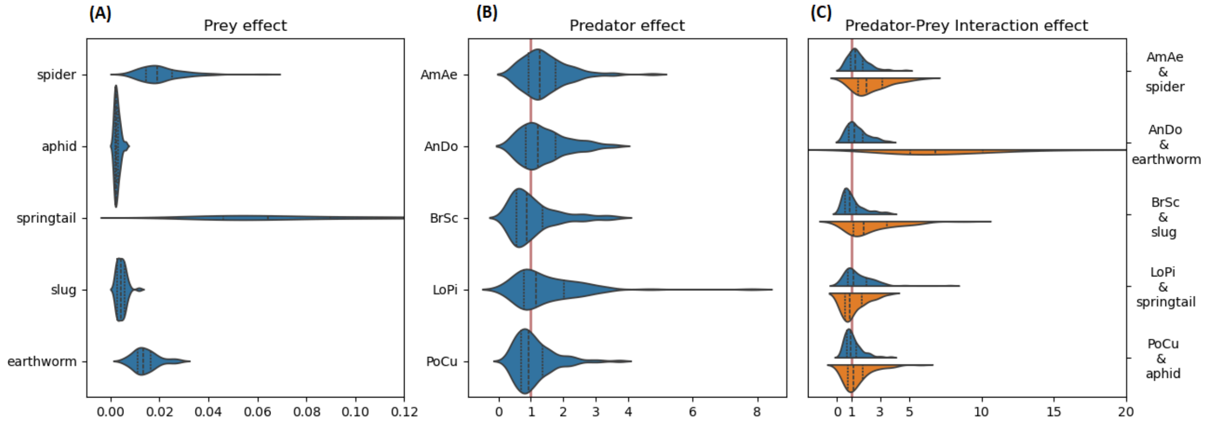


**FIGURE 15** - *Analysis of parameter estimates derived from the decomposition of predation rate:* $\lambda^{p,c} = e^{\alpha_{\lambda}^{p}} \cdot e^{\delta_{\lambda}^{c}} \cdot e^{\gamma_{\lambda}^{p,c}}$ *(see M&M – parameter decomposition). Panel* ***(A)*** *shows* $e^{\alpha_{\lambda}^{p}}$*, the exponentials of prey effects estimate for the five types of prey considered. Panel* ***(B)*** *shows* $e^{\delta_{\lambda}^{c}}$*, the exponentials of predator effects estimate for five predator species selected from the 25 studied. Panel* ***(C)*** *shows* $e^{\gamma_{\lambda}^{p,c}}$ *, the exponentials of prey-predator interaction effects for 5 emblematic interactions.*

In **Figure 15 (A)**, we observe that prey effects are by far the most contrasted. With a mean effect ranging from 0.005 for aphids to 0.05 for springtails, it is by far the primary factor of variability in predation rate. This marked contrast can be explained by the great diversity of prey types considered, ranging from very abundant small prey like springtails to much larger prey like earthworms, and/or rarer prey like slugs. The characteristics of predation on each prey, especially the number of predation events, could only be very different.

Interaction effects seem to be secondary factors of variability in predation rate estimates (**Figure 15 (C)**). Although they are of the same order of magnitude as predator effects, they show real differences among predator-prey couples allowing the identification of specialization effects. In particular, the interaction effect of *A. Dorsalis* on earthworms is the most significant interaction effect (6 times greater than the average interaction effect), which is consistent with the literature. Conversely, the interaction effect of *L.Pilicornis* on springtails does not reflect its specialization, which is probably due to *L.Pilicornis* not significantly differing from other carabid species, neither in field data (average frequency of individuals tested positive – see Supp. Mat. 4) nor in laboratory data (average voracity and average digestion speed).

In contrast (**Figure 15 (B)**), carabid effects are all very similar. This observation leads us to believe either that carabid species has no effect on predation rate, or that information on voracity is carried by other parameters within the model. In particular, the voracity of carabid beetles under controlled conditions is already considered in the digestion properties estimates, and could be redundant with the information carried by the carabid beetle effect. Information on species voracity could also be carried directly by interaction effects estimates, which are slightly correlated with carabid effects.

**Supplementary Material 8 |** Historical models' predictions

---

All frequentist models presented in **Figure 16** are in the form: $\lambda = p\frac{R}{D}$

$\lambda$: Predation rate
$p$: Proportion of predators tested positive
$R$: Number of preys eaten when tested positive

$$R = 1 \; or \; \frac{-ln(1-p)}{p} \; or \; \frac{(1-k)}{p} \, ln\left(\frac{1-k}{1-p-k}\right)$$

According respectfully to [Dempster 1960], to [Nakamura et Nakamura 1977, Greenstone 1979] using Poisson distribution to count the number of preys eaten, and [Lister 1987 (a)] with a proportion of inactive predators $k = 0.5$.

$D$: Measure of molecular signal decay: $D = D_{mean} \; or \; D_{max} \; or \; H_{50}$, with $D_{mean}$ and $D_{max}$ the mean and maximum digestion time after which the prey can no longer be detected, and $H_{50} = -\frac{\beta_1}{\beta_0}$ the detectability half-life of the prey according to our equation of prey decay (see Table 1 in Supp. Mat. 1 for the details on $\beta_1$ and $\beta_0$).

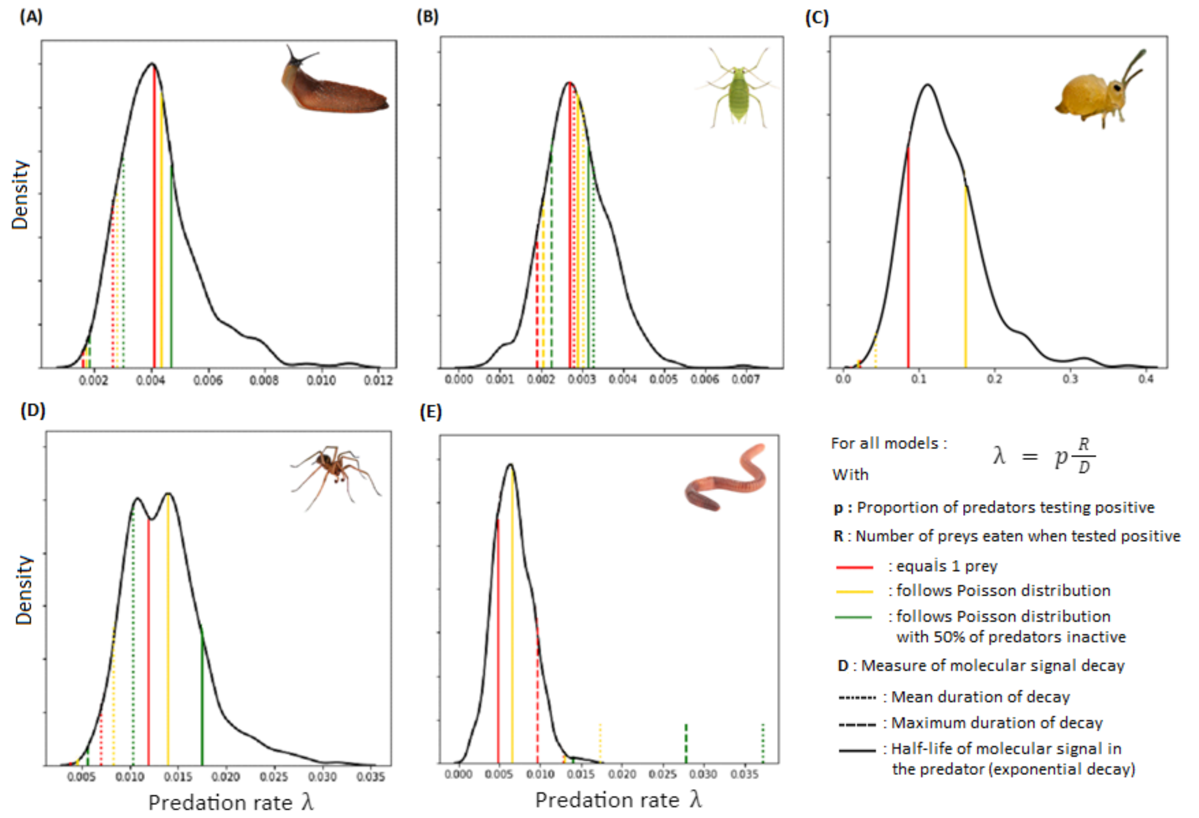


**FIGURE 16 -** ***Comparison of predation rates (per hour) of Poecilus cupreus on 5 types of prey, estimated using different methods****. The thick black line represents the posterior distribution of predation rate λ obtained with our model. Vertical lines are the predation rate based on previously published models. They differ in the way they calculate the number of prey consumed, which can be*

*fixed at 1 [Dempster 1960], estimated from a Poisson distribution [Nakamura and Nakamura 1977, Greenstone 1979], or take into account the proportion of inactive predators [Lister et al. 1987]. They also differ in the way digestion (i.e. degradation of the molecular signal) is accounted for, between a simple (average or maximal) duration ($D = D_{mean}$ or $D_{max}$) [Dempster 1960, Rothschild 1966, Nakamura and Nakamura 1977, Greenstone 1979, Kuperstein 1974 & 1979, Sunderland 1987, Lister 1987a] or a half-life underlying an exponential decay model [Lister 1987b, Sopp et al. 1992, Naranjo and Hagler 2000, Greenstone et al. 2010, 2014, Andow and Paula 2023].*

The HBM method provides estimates that are consistent with the ones provided by previously published models on our application example. The main source of divergence among published models estimates appears to lie in the way in which molecular signal decay is accounted for. Unsurprisingly [Naranjo and Hagler 2000], the use of detectability periods (average or maximum $D_{mean}$ or $D_{max}$) of the molecular signal in the predator leads to a strong underestimation of predation rate, whereas the use of half-lives leads to predictions more in line with our estimates. This confirms the relevance of using half-lives as recommended by [Greenstone et al. 2010, 2014]. Conversely, consideration of the number of preys eaten when tested positive, for example in the form of a Poisson distribution [Nakamura and Nakamura 1977, Greenstone 1979], appears to be relatively secondary. This may be due to the fact that predation rates are moderate in our application example. Indeed, as the probability of two events being detected by the same test increases, the gap between predictions with and without considering the number of prey eaten increases (e.g. with Poecilus cupreus on springtails). Finally, defining a proportion of the captured population as inactive [Lister et al. 1987], leads to an overestimation of the predation rates compared to other models and our estimates. This is consistent with the fact that, since they were caught with Barber traps, all individuals in our dataset are active.

**Supplementary Material 9** |Barber Trap Correction

Our model assumes that the end of the predation process occurs when the carabid falls into the Barber trap. The digestion process however continues until the Barber trap is collected, which results in a time lag between the end of predation and the end of digestion. As the trap were collected every 12 hours, the duration of the time lag varies between 0 and 12 hours, but its exact value remains unknown. If ignored, this time lag may lead to an underestimation of predation rate. The second version of our model takes into account the time lag using random or average residence time (6h) in the trap. Model calibration was performed on both and the MCMC algorithm converged well in both cases. This allows us to confirm that our model remains identifiable even when predation is not constant. These encouraging results invite us to attempt more substantial variations of predation rate, possibly supported by data on predator traits (e.g. nycthemeral rhythm).

Samples and estimates for both random and average versions are given with the Code associated with the present study.

**Supplementary Material 10 |** Carabid Codes

| SpeciesCode | SpeciesName |
|---|---|
| AbOv | Abax ovalis |
| AbPa | Abax parallelepipedus |
| AbPar | Abax parallelus |
| AcEl | Acupalpus elegans |
| AcMe | Acupalpus meridianus |
| AcSp | Acupalpus sp |
| AgAf | Agonum afrum |
| AgDu | Agonum duftschmidi |
| AgGr | Agonum gracilipes |
| AgMu | Agonum muelleri |
| AgNi | Agonum nigrum |
| AgSe | Agonum sexpunctatum |
| AgSp | Agonum sp |
| AgVi | Agonum viduum |
| AmAe | Amara aenea |
| AmCo | Amara convexior |
| AmCu | Amara curta |
| AmCur | Amara cursitans |
| AmEu | Amara eurynota |
| AmFa | Amara familiaris |
| AmOv | Amara ovata |
| AmPl | Amara plebeja |
| AmSi | Amara similata |
| AmSp | Amara sp |
| AmbNi | Amblystomus niger |
| AmbSp | Amblystomus sp |
| AnDo | Anchomenus dorsalis |
| AnBi | Anisodactylus binotatus |
| ApBo | Aptinus bombarda |
| AsFl | Asaphidion gr. flavipes |
| BaBu | Badister bullatus |
| BaSo | Badister sodalis |
| BaSp | Badister sp |
| BeQu | Bembidion quadrimaculatum |
| BeSp | Bembidion sp |
| BlMu | Blethisa multipunctata |
| BrCr | Brachinus crepitans |
| BrEx | Brachinus explodens |
| BrSc | Brachinus sclopeta |
| BrSp | Brachinus sp |

| | |
|---|---|
| BraCa | Bradycellus caucasicus |
| BraHa | Bradycellus harpalinus |
| CalEr | Calathus erratus |
| CalFu | Calathus fuscipes |
| CalMe | Calathus melanocephalus |
| CalMet | Calathus metallicus |
| CalMi | Calathus micropterus |
| CalMo | Calathus mollis |
| CalSp | Calathus sp |
| CaoIn | Calosoma inquisitor |
| CaArv | Carabus arcensis |
| CaAr | Carabus auronitens |
| CaAu | Carabus auratus |
| CaCa | Carabus cancellatus |
| CaCo | Carabus convexus |
| CaCor | Carabus coriaceus |
| CaFr | Carabus fabricii |
| CaGl | Carabus glabratus |
| CaGr | Carabus granulatus |
| CaHo | Carabus hortensis |
| CaIr | Carabus irregularis |
| CaLi | Carabus linnei |
| CaMn | Carabus monilis |
| CaNe | Carabus nemoralis |
| CaOb | Carabus obsoletus |
| CaPr | Carabus problematicus |
| CaSc | Carabus scheidleri |
| CaSp | Carabus sp |
| CaSpl | Carabus splendens |
| CaSy | Carabus sylvestris |
| CaUl | Carabus ulrichii |
| CaVi | Carabus violaceus |
| ChTr | Chlaenius tristis |
| Cica | Cicindela campestris |
| ClFo | Clivina gr. fossor |
| CyAt | Cychrus attenuatus |
| CyCa | Cychrus caraboides |
| CyGe | Cylindera germanica |
| CyAx | Cymindis axillaris |
| CyHu | Cymindis humeralis |
| DeTa | Deltomerus tatricus |
| DeAt | Demetrias atricapillus |
| DiGe | Diachromus germanus |
| DiSp | Diachromus sp |
| DrFe | Dromius fenestratus |
| DrDe | Drypta dentata |

| | |
|---|---|
| EpSe | Epaphius secalis |
| HaAf | Harpalus affinis |
| HaAt | Harpalus atratus |
| HaCa | Harpalus calceatus |
| HaCu | Harpalus cupreus |
| HaDm | Harpalus dimidiatus |
| HaDi | Harpalus distinguendus |
| HaHo | Harpalus honestus |
| HaLa | Harpalus latus |
| HaLu | Harpalus luteicornis |
| HaPu | Harpalus punctulatus |
| HaQu | Harpalus quadripunctatus |
| HaRu | Harpalus rubripes |
| HaRuf | Harpalus rufipalpis |
| HaSi | Harpalus signaticornis |
| HaSm | Harpalus smaragdinus |
| HaSp | Harpalus sp |
| HaTa | Harpalus tardus |
| HaTe | Harpalus tenebrosus |
| LeFe | Leistus ferrugineus |
| LePi | Leistus piceus |
| LeRuf | Leistus rufomarginatus |
| LiHo | Licinus hoffmannseggi |
| LoPi | Loricera pilicornis |
| MeLa | Metallina lampros |
| MePr | Metallina properans |
| MeSp | Metallina sp |
| MiMa | Microlestes maurus |
| MiMi | Microlestes minutulus |
| MiSp | Microlestes sp |
| MoEl | Molops elatus |
| MoPi | Molops piceus |
| NeBr | Nebria brevicollis |
| NeSa | Nebria salina |
| NeSp | Nebria sp |
| NeTa | Nebria tatrica |
| NoBi | Notiophilus biguttatus |
| NoPa | Notiophilus palustris |
| NoQu | Notiophilus quadripunctatus |
| NoRu | Notiophilus rufipes |
| NoSp | Notiophilus sp |
| nous | Notiophilus substriatus |
| OcDe | Ocydromus deletus |
| OcGe | Ocydromus genei |
| OcTe | Ocydromus tetracolus |
| OpAz | Ophonus azureus |

| | |
|---|---|
| OpPu | Ophonus gr. puncticeps |
| OpRu | Ophonus rufibarbis |
| OpSu | Ophonus subquadratus |
| OpSp | Ophonus sp |
| PaBi | Paratachys bistriatus |
| PaSp | Paratachys sp |
| PaMe | Parophonus mendax |
| PhiBi | Philochthus biguttatus |
| PhiIr | Philochthus iricolor |
| PhiLu | Philochthus lunulatus |
| PhiSp | Philochthus sp |
| PhOb | Phyla obtusa |
| PhSp | Phyla sp |
| PlAs | Platynus assimilis |
| PoCu | Poecilus cupreus |
| PoKu | Poecilus kugelanni |
| PoLe | Poecilus lepidus |
| PoSe | Poecilus sericeus |
| PoSp | Poecilus sp |
| PoVe | Poecilus cf. versicolor |
| PoCo | Polistichus connexus |
| PsRu | Pseudoophonus rufipes |
| PtAe | Pterostichus aethiops |
| PtAn | Pterostichus anthracinus |
| PtBl | Pterostichus blandulus |
| PtBu | Pterostichus burmeisteri |
| PtCo | Pterostichus cordatus |
| PtFo | Pterostichus foveolatus |
| PtMa | Pterostichus madidus |
| PtMe | Pterostichus melanarius |
| PtMel | Pterostichus melas |
| PtMo | Pterostichus morio |
| PtNi | Pterostichus niger |
| PtNig | Pterostichus nigrita |
| PtOb | Pterostichus oblongopunctatus |
| PtOv | Pterostichus ovoideus |
| PtPi | Pterostichus pilosus |
| PtPu | Pterostichus pumilio |
| PtSp | Pterostichus sp |
| PtSt | Pterostichus strenuus |
| PtUn | Pterostichus unctulatus |
| PtVe | Pterostichus vernalis |
| StMi | Stenolophus mixtus |
| StSp | Stenolophus sp |
| StTe | Stenolophus teutonus |
| StPu | Stomis pumicatus |

| | |
|---|---|
| SyFo | Syntomus foveatus |
| SySp | Syntomus sp |
| SyOb | Syntomus obscuroguttatus |
| SyTr | Syntomus truncatellus |
| SynVi | Synuchus vivalis |
| TrLa | Trechus latus |
| TrPi | Trechus pilisensis |
| TrPu | Trechus pulchellus |
| TrQu | Trechus quadristriatus |
| TrRu | Trechus rubens |
| TrSp | Trechus sp |
| TrSt | Trechus striatulus |
| TriLa | Trichotichnus laevicollis |
| ZaTe | Zabrus tenebrioides |

**Supplementary Material 11 |** Community structure throughout the cropping season

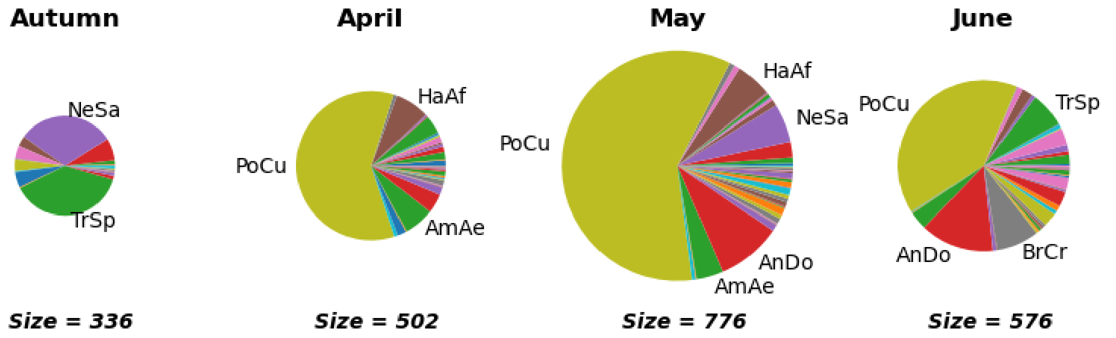


**FIGURE 6 - Changes in carabid community structure (composition and activity-density) during the cropping season -** *For each sampling session the activity-density (circle radius) and the proportion of each carabid species in the community (circle division) are depicted. Data from wheat fields in 4 French regions were pooled (see Supp. Mat. 10 for carabid code names)*

The carabid communities in autumn were the smallest in size and the least diverse in species, with a total of 336 individuals caught of 20 species, compared with an average of 618 individuals of 35, 40, and 41 species for the spring sessions. Moreover, species of the *Nebria* and *Trechus* genus accounted for more than 75% of individuals caught in autumn, but less than 10% in spring communities. From April to June, variations were weaker. The main differences concerned global activity-density, and the proportion of *A. dorsalis*, which increases throughout the spring, *B. crepitans*, that explodes in June, and *Harpalus affinis*, which drops in June.
