## Supplementary material for "Unveiling the Hidden Feast: from molecular detection to predation rate – An example on biological control by generalist predators": Main Code

### *Unveiling the Hidden Feast: translating molecular detection into predation rate – An example on biological control by generalist predators in agricultural fields.*

#### *Code and documentation*

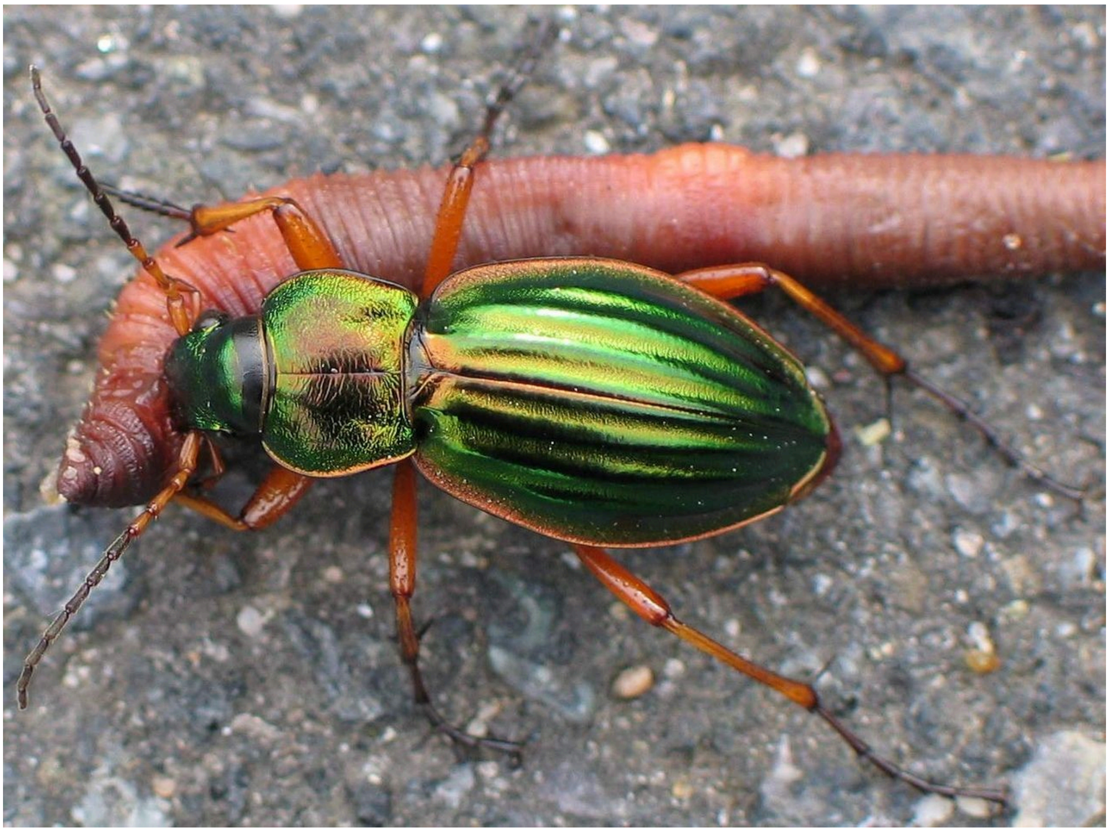

##### IMPORTING USEFUL LIBRARIES

⚠ *The versions of Python and the different libraries used in this script must strictly match the versions listed in the Watermark section at the end of the document* ⚠

In [3]:
```
import sys
import platform
import scipy
import datetime
import pandas as pd
import pymc as pm
import numpy as np
import matplotlib
import matplotlib.pyplot as plt
import seaborn as sns
import arviz as az
from scipy.special import logit
import pytensor
import pytensor.tensor as pt
import xarray as xr
import warnings

from scipy.io import netcdf
```

```
from Data.Utils import *
from Data.Data2 import *
from Figures.Figures import *
```

```
Running on PyMC v5.6.1
```

In [4]:
```
%config InlineBackend.figure_format = 'retina'
az.style.use("arviz-darkgrid")
rng = np.random.default_rng(42)
warnings.filterwarnings('ignore')
```

##### IMPORTING AND PREPROCESSING DATA

The laboratory and field data are stored as Excel spreadsheets in the /Data folder. The details of the import and pre-processing steps are provided in the Data.py script, which in particular defines the Data class. In the remainder of this document, the laboratory and field data are separated into *df_dig* and *df_pred* dataframes, whose columns and first rows are shown below.

In [6]:
```
# Importing and pre-processing data

dataset = Dataset(path_to_fig=os.getcwd(),
                  import_ = True,
                  preprocess_ = True)

# Definition of df_dig and df_pred

df_dig = dataset.get_data(type_de_données="labo")

df_pred = dataset.get_data(type_de_données="terrain",
                           liste_variables=["Carabe", "Proie", "Couple", "Detect
                           liste_carabes= list(pd.unique(df_dig.Carabe)))
```

In [7]:
```
df_dig.head(5)
```

Out[7]:

| | Couple | Carabe | Temps de Digestion | Proie Consommée | Detection |
|---|---|---|---|---|---|
| **0** | PoCu_slug | PoCu | 3.0 | slug | 1 |
| **1** | PoCu_spider | PoCu | 24.0 | spider | 0 |
| **2** | PoCu_springtail | PoCu | 3.0 | springtail | 0 |
| **3** | AmAe_aphid | AmAe | 24.0 | aphid | 0 |
| **4** | AmAe_spider | AmAe | 3.0 | spider | 0 |

In [8]:
```
df_pred.head(5)
```

Out[8]:

| | Carabe | Proie | Couple | Detection |
|---|---|---|---|---|
| **0** | AcMe | earthworm | AcMe_earthworm | 0 |
| **1** | AcMe | earthworm | AcMe_earthworm | 0 |
| **2** | AcMe | earthworm | AcMe_earthworm | 0 |
| **3** | AcMe | earthworm | AcMe_earthworm | 0 |
| **4** | AcMe | earthworm | AcMe_earthworm | 0 |

##### MODEL DEFINITION

The hierarchical Bayesian model used in our study is defined in the cell below. Several other versions of it have been developed (digestion model, model with correction for the delay in the Barber trap, for example), but they are not presented here for the sake of clarity.

In [10]:
```
def Model_Digestion_et_Predation(df_dig, df_pred):

    """
    Parameters
    ----------

    df_dig : Dataframe
        Data from the digestion experiment under controlled conditions.
    df_pred : Dataframe
        Data on prey detection in the gut contents of predators caught in in nat

    Returns
    -------
    model : pyMC Model Object
        A pyMC Model ready for training.
    """

    assert all([(proie in pd.unique(df_dig["Proie Consommée"]))
                for proie in pd.unique(df_pred["Proie"])]), "prey type list diff
    assert all([(carabe in pd.unique(df_dig["Carabe"]))
                for carabe in pd.unique(df_pred["Carabe"])]), "predator list dif

    if len(pd.unique(df_dig["Couple"])) != len(pd.unique(df_dig["Carabe"]))*len(

        for proie in pd.unique(df_dig["Proie Consommée"]) :
            for carabe in pd.unique(df_dig["Carabe"]) :

                if not ("_".join([carabe, proie]) in pd.unique(df_dig["Couple"])

                    couple_row = {'Couple': "_".join([carabe, proie]),
                                  'Carabe': carabe,
                                  'Temps de Digestion': 0,
                                  'Proie Consommée' : proie,
                                  'Detection' : 1}

                    df_dig.loc[len(df_dig)] = couple_row

    observations_prédation_idx = np.asarray(df_pred.index)
    observations_digestion_idx = np.asarray(df_dig.index)
```

```
time_idx, times = pd.factorize(df_dig["Temps de Digestion"], sort=True)

proies_digestion_idx, proies = pd.factorize(df_dig["Proie Consommée"], sort=
proies_prédation_idx, _ = pd.factorize(df_pred["Proie"], sort=True)

carabes_digestion_idx, carabes = pd.factorize(df_dig["Carabe"], sort=True)
carabes_prédation_idx, _ = pd.factorize(df_pred["Carabe"], sort=True)

couples_digestion_idx, couples = pd.factorize(df_dig["Couple"], sort=True)
couples_prédation_idx, _ = pd.factorize(df_pred["Couple"], sort=True)

proies_de_couple_idx = np.asarray(
    [list(proies).index(couple.split("_")[1]) for couple in couples])
carabes_de_couple_idx = np.asarray(
    [list(carabes).index(couple.split("_")[0]) for couple in couples])
couples_de_couple_idx = np.asarray([list(couples).index(
    couple) for couple in couples])

coords = {"observations_prédation": observations_prédation_idx,
          "observations_digestion": observations_digestion_idx,
          "proies": proies,
          "carabes": carabes,
          "couples": couples}

# DEFINITION DU MODELE
# ---------------------------------------------------------------------------

with pm.Model(coords=coords) as model:

    # Conversion des variables observées au format np.array
    test_prédation_obs = pm.ConstantData("test_prédation_obs", df_pred["Dete
                                         dims="observations_prédation")
    test_digestion_obs = pm.ConstantData("test_obs", df_dig["Detection"],
                                         dims="observations_digestion")
    temps_digestion = pm.ConstantData("temps_digestion", df_dig["Temps de Di
                                      dims="observations_digestion")

    # HYPERPRIORS

    # On the intercept beta0 (intercept mu_beta0, prey effect alpha_beta0_p,
    # and the interaction effect gamma_beta0_pc)

    # prey effect
    alpha_beta0_p = pm.Normal("alpha_beta0_p", mu=0, tau=1, dims="proies")

    # predator effect
    mu_delta_beta0_c = pm.Normal("mu_delta_beta0_c", mu=0, tau=100)
    log_sigma_delta_beta0_c = pm.Uniform("log_sigma_delta_beta0_c", 0, 1)
    tau_delta_beta0_c = 1/np.exp(log_sigma_delta_beta0_c)

    # prey-predator interaction effect
    mu_gamma_beta0_pc = pm.Normal("mu_gamma_beta0_pc", mu=0, tau=100)
    log_sigma_gamma_beta0_pc = pm.Uniform("log_sigma_gamma_beta0_pc", 0, 1)
    tau_gamma_beta0_pc = 1/np.exp(log_sigma_gamma_beta0_pc)

    # On the slope of the digestion curve beta1 (mean value mu_beta1, and va
    mu_log_beta1 = pm.Normal("mu_log_beta1", mu=-2, tau=100)
    sigma_log_beta1 = pm.Uniform("sigma_log_beta1", 0, 1)
    tau_log_beta1 = 1/np.exp(sigma_log_beta1)
```

```
        # On the log of the predation rate lambda (intercept mu_loglambda, prey
        # and the interaction effect gamma_loglambda_pc)

        # prey effect
        alpha_loglambda_p = pm.Normal("alpha_loglambda_p", mu=0, tau=0.05, dims=

        # predator effect
        mu_delta_loglambda_c = pm.Normal("mu_delta_loglambda_c", mu=0, tau=100)
        log_sigma_delta_loglambda_c = pm.Uniform("log_sigma_delta_loglambda_c",
        tau_delta_loglambda_c = 1/np.exp(log_sigma_delta_loglambda_c)

        # prey-predator interaction effect
        mu_gamma_loglambda_pc = pm.Normal("mu_gamma_loglambda_pc", mu=0, tau=100
        log_sigma_gamma_loglambda_pc = pm.Uniform("log_sigma_gamma_loglambda_pc"
        tau_gamma_loglambda_pc = 1/np.exp(log_sigma_gamma_loglambda_pc)

        # PRIORS

        # On the parameters of the digestion curve
        delta_beta0_c = pm.Normal("delta_beta0_c", mu=mu_delta_beta0_c,
                                  tau=tau_delta_beta0_c, dims="carabes")

        gamma_beta0_pc = pm.Normal("gamma_beta0_pc", mu=mu_gamma_beta0_pc,
                                   tau=tau_gamma_beta0_pc, dims="couples")

        # On predation rate
        delta_loglambda_c = pm.Normal("delta_loglambda_c", mu=mu_delta_loglambda
                                      tau=tau_delta_loglambda_c, dims="carabes")

        gamma_loglambda_pc = pm.Normal("gamma_loglambda_pc", mu=mu_gamma_loglamb
                                       tau=tau_gamma_loglambda_pc, dims="couples

        # MODEL

        # Digestion

        log_beta1_pc = pm.Normal("log_beta1_pc", mu=mu_log_beta1,
                                 tau=tau_log_beta1, dims="couples")

        beta1_pc = pm.Deterministic("beta1_pc", (-1)*np.exp(log_beta1_pc),
                                    dims="couples")

        beta0_pc = pm.Deterministic("beta0_pc", alpha_beta0_p[proies_de_couple_i
                                   + delta_beta0_c[carabes_de_couple_idx]
                                   + gamma_beta0_pc[couples_de_couple_idx],
                                   dims="couples")

        Pi_t = pm.Deterministic("Pi_t", beta0_pc[couples_digestion_idx]
                               + beta1_pc[couples_digestion_idx] *
                               temps_digestion,
                               dims="observations_digestion")  # Probability of

        # Predation

        lambda_pc = pm.Deterministic("lambda_pc", np.exp(alpha_loglambda_p[proie
                                                    + delta_loglambda_c[car
                                                    + gamma_loglambda_pc[cc
                                     dims="couples")  # Predation rate (hourly)
```

```
        # Digestion and predation combination

        T0 = 0 ; Tmax = 2000

        I_pc = pm.Deterministic("I_pc", (1/beta1_pc)*(np.log(1 + np.exp(beta0_pc
                                                      - np.log(1 + np.exp(beta0_
                                dims="couples")  # Integral of the digestion cur

        pI = pm.Deterministic("pI", 1 - pm.math.maximum(0.001,
                                                  np.exp(-lambda_pc[couples_prédati
                                                         I_pc[couples_prédation_idx
                              dims="observations_prédation")  # Probability of p

        # OBSERVATION PROCESS

        test_prédation_pred = pm.Bernoulli("test_prédation_pred", p=pI, observed
                                            dims="observations_prédation")
        test_digestion_pred = pm.Bernoulli("test_dig_pred", logit_p=Pi_t, observ
                                            dims="observations_digestion")

    return model
```

#### IMPLEMENTATION AND VISUALIZATION OF THE MODEL

In the following cell, we construct an instance of the model by implementing the definition function described above, using the available laboratory and field datasets as arguments (df_dig and df_pred).

We also provide a visualization of the model (i.e., its various variables, their dimensions, and the links between them), automatically generated from its definition using the Graphviz module.

In [12]:
```
model = Model_Digestion_et_Predation(df_dig, df_pred)

Graph = pm.model_to_graphviz(model)
Graph.view()
Graph
```

Out[12]:

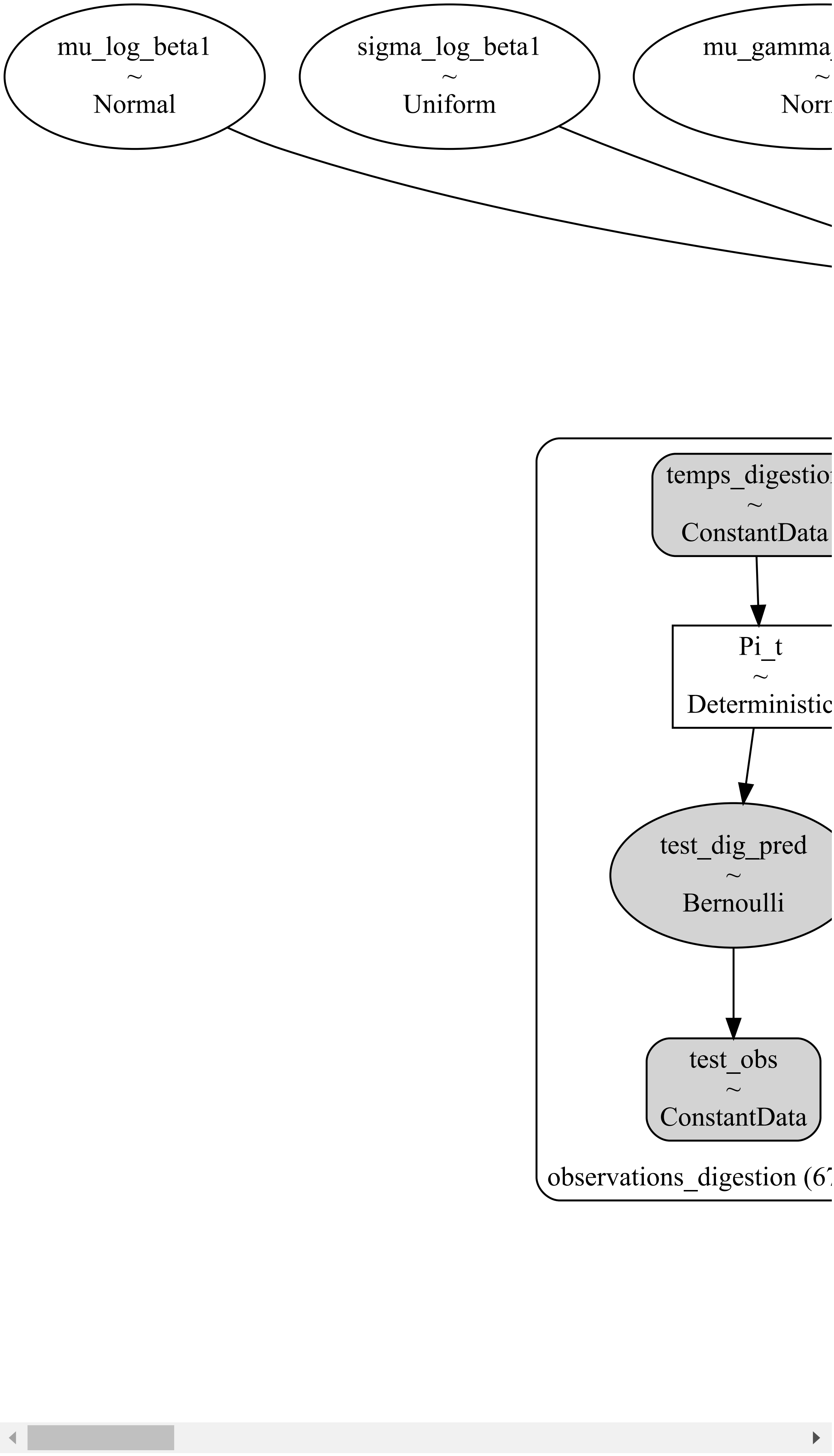
mu_log_beta1
~
Normal
sigma_log_beta1
~
Uniform
temps_digestio
~
ConstantData
Pi_t
~
Deterministic
test_dig_pred
~
Bernoulli
test_obs
~
ConstantData
observations_digestion (6

#### MODEL CALIBRATION

The sample() function is used to calibrate the model, i.e. to run markov chain simulations to obtain the posterior distributions of its parameters. In the cell below it is given as an example, with its various arguments (sample size, number of chains, burnin, etc.) but as calibration can be excessively long, it is preferable to retrieve samples that have already been simulated directly from the /Samples folder.

*Path_to_InferenceData* and *InferenceData* indicate the location and the name of the sample. These two fields can be modified to load the appropriate sample, depending on the model version and the type of calibration one wishes to study. For example, certain samples have been obtained using only the data from one particular capture session.

In [14]:
```
Path_to_InferenceData = './Samples/Echantillons_Modèle_Final/'
InferenceData = 'Echantillon_Model_Digestion_et_Predation_Informative_Prior_27sp
```

In [15]:
```
try :

    DataInference = xr.open_dataset("".join([Path_to_InferenceData, InferenceDat
    Echantillons = az.convert_to_inference_data(DataInference)

except :

    InferenceData_tostock = InferenceData[:-3] + "_tostock.nc"

    Echantillons = pm.sample(draws=3000,
                             chains=4,
                             tune=1000,
                             progressbar=True,
                             model=model,
                             compute_convergence_checks=True)

    DataSet_Echantillons = az.convert_to_dataset(Echantillons)
    DataSet_Echantillons.to_netcdf("".join([Path_to_InferenceData, InferenceData
```

#### CONVERGENCE TESTS

Convergence tests are used to test the quality of the samples, i.e. that the three or four Markov chains that make up the sample underlie the same *posterior* distribution, i.e. that the chains converge.

Many random variables are defined during the model implementation, and not all of them are necessarily relevant for display/analysis. A list of the names of relevant variables can be provided as input to the functions below for clearer presentation of the results.

In [18]:
```
var_names = ["alpha_beta0_p", "mu_delta_beta0_c", "log_sigma_delta_beta0_c", "mu
             "sigma_log_beta1", "alpha_loglambda_p", "mu_delta_loglambda_c", "log
             "log_sigma_gamma_loglambda_pc", "delta_beta0_c", "gamma_beta0_pc", "

var_names_summary = ['mu_delta_beta0_c', 'log_sigma_delta_beta0_c', 'mu_gamma_be
                     'mu_log_beta1', 'sigma_log_beta1', 'mu_delta_loglambda_c',
                     'log_sigma_gamma_loglambda_pc']
```

*Calculation and display of convergence metrics*

The most important convergence metric calculated by the summary() function from the pyMC library based on the sample is the Gelman-Rubin statistic, also known as r_hat (see table below). This statistic measures the similarity between the three simulated Markov chains. The closer it is to 1, the more similar the Markov chains are. Conversely, if it exceeds 1.05, we can consider that at least one Markov chain has not converged, and that the sample does not accurately represent the *posterior* distribution.

In [21]:
```
try :

    print(summary.loc[var_names_summary])

except :

    summary = az.summary(Echantillons)
    print(summary.loc[var_names_summary])
```

```
                              mean     sd  hdi_3%  hdi_97%  mcse_mean  \
mu_delta_beta0_c             0.060  0.098  -0.126    0.241      0.001   
log_sigma_delta_beta0_c      0.354  0.256   0.000    0.825      0.002   
mu_gamma_beta0_pc            0.061  0.098  -0.121    0.248      0.001   
log_sigma_gamma_beta0_pc     0.168  0.149   0.000    0.447      0.002   
mu_log_beta1                -2.106  0.087  -2.267   -1.943      0.001   
sigma_log_beta1              0.082  0.076   0.000    0.223      0.001   
mu_delta_loglambda_c        -0.010  0.101  -0.202    0.177      0.001   
log_sigma_delta_loglambda_c  0.207  0.188   0.000    0.560      0.001   
mu_gamma_loglambda_pc       -0.012  0.101  -0.201    0.179      0.001   
log_sigma_gamma_loglambda_pc  0.094  0.088   0.000    0.257      0.001   

                              mcse_sd  ess_bulk  ess_tail  r_hat  
mu_delta_beta0_c                0.001   24897.0   15391.0    1.0  
log_sigma_delta_beta0_c         0.002   14422.0   13798.0    1.0  
mu_gamma_beta0_pc               0.001   15521.0   14261.0    1.0  
log_sigma_gamma_beta0_pc        0.001    7096.0    8067.0    1.0  
mu_log_beta1                    0.000   18627.0   15724.0    1.0  
sigma_log_beta1                 0.000   13816.0   10027.0    1.0  
mu_delta_loglambda_c            0.001   18452.0   15157.0    1.0  
log_sigma_delta_loglambda_c     0.001   13855.0    9844.0    1.0  
mu_gamma_loglambda_pc           0.001   12141.0   14334.0    1.0  
log_sigma_gamma_loglambda_pc    0.001   11993.0    9854.0    1.0  
```

*Visualization of chains for graphical verification of convergence*

In the cell below, the trace plots for the graphical validation of MCMC convergence. They display the raw Markov Chains (right panels) and posterior distributions (left panels) of the most important parameters of the model. On each panel, several parameters can be displayed (e.g. the five prey effects). It that case, they are discriminated using a color code

In [24]:
```
az.plot_trace(Echantillons,
              var_names = var_names,
              divergences = "top",
              figsize = (0.5*len(var_names),2*len(var_names)),
              combined=False)
```

```
Out[24]: array([[<Axes: title={'center': 'alpha_beta0_p'}>,
                 <Axes: title={'center': 'alpha_beta0_p'}>],
                [<Axes: title={'center': 'mu_delta_beta0_c'}>,
                 <Axes: title={'center': 'mu_delta_beta0_c'}>],
                [<Axes: title={'center': 'log_sigma_delta_beta0_c'}>,
                 <Axes: title={'center': 'log_sigma_delta_beta0_c'}>],
                [<Axes: title={'center': 'mu_gamma_beta0_pc'}>,
                 <Axes: title={'center': 'mu_gamma_beta0_pc'}>],
                [<Axes: title={'center': 'log_sigma_gamma_beta0_pc'}>,
                 <Axes: title={'center': 'log_sigma_gamma_beta0_pc'}>],
                [<Axes: title={'center': 'mu_log_beta1'}>,
                 <Axes: title={'center': 'mu_log_beta1'}>],
                [<Axes: title={'center': 'sigma_log_beta1'}>,
                 <Axes: title={'center': 'sigma_log_beta1'}>],
                [<Axes: title={'center': 'alpha_loglambda_p'}>,
                 <Axes: title={'center': 'alpha_loglambda_p'}>],
                [<Axes: title={'center': 'mu_delta_loglambda_c'}>,
                 <Axes: title={'center': 'mu_delta_loglambda_c'}>],
                [<Axes: title={'center': 'log_sigma_delta_loglambda_c'}>,
                 <Axes: title={'center': 'log_sigma_delta_loglambda_c'}>],
                [<Axes: title={'center': 'mu_gamma_loglambda_pc'}>,
                 <Axes: title={'center': 'mu_gamma_loglambda_pc'}>],
                [<Axes: title={'center': 'log_sigma_gamma_loglambda_pc'}>,
                 <Axes: title={'center': 'log_sigma_gamma_loglambda_pc'}>],
                [<Axes: title={'center': 'delta_beta0_c'}>,
                 <Axes: title={'center': 'delta_beta0_c'}>],
                [<Axes: title={'center': 'gamma_beta0_pc'}>,
                 <Axes: title={'center': 'gamma_beta0_pc'}>],
                [<Axes: title={'center': 'delta_loglambda_c'}>,
                 <Axes: title={'center': 'delta_loglambda_c'}>],
                [<Axes: title={'center': 'gamma_loglambda_pc'}>,
                 <Axes: title={'center': 'gamma_loglambda_pc'}>],
                [<Axes: title={'center': 'beta1_pc'}>,
                 <Axes: title={'center': 'beta1_pc'}>]], dtype=object)
```

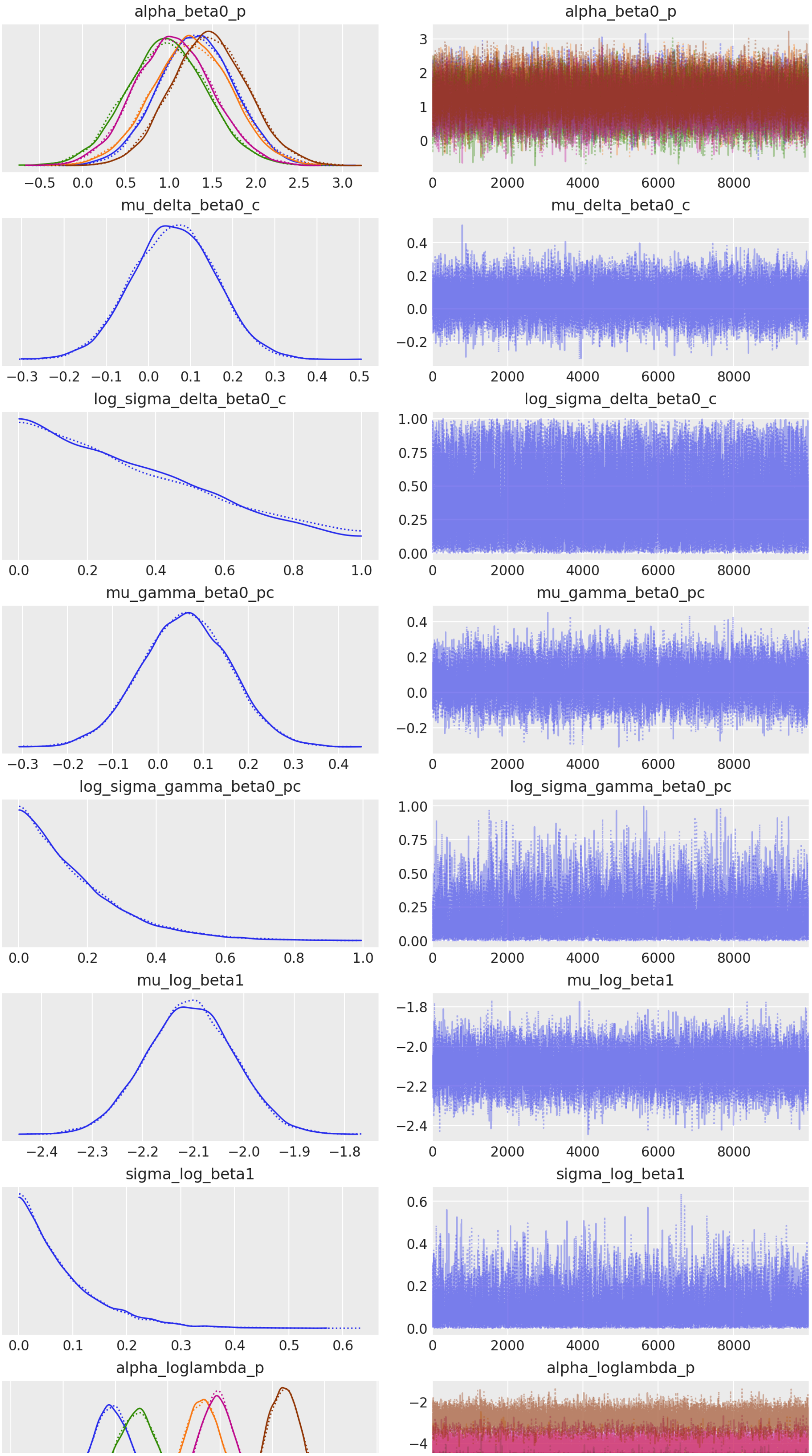
alpha_beta0_p
−0.5 0.0 0.5 1.0 1.5 2.0 2.5 3.0
alpha_beta0_p
3 2 1 0
0 2000 4000 6000 8000
mu_delta_beta0_c
−0.3 −0.2 −0.1 0.0 0.1 0.2 0.3 0.4 0.5
mu_delta_beta0_c
0.4 0.2 0.0 −0.2
0 2000 4000 6000 8000
log_sigma_delta_beta0_c
0.0 0.2 0.4 0.6 0.8 1.0
log_sigma_delta_beta0_c
1.00 0.75 0.50 0.25 0.00
0 2000 4000 6000 8000
mu_gamma_beta0_pc
−0.3 −0.2 −0.1 0.0 0.1 0.2 0.3 0.4
mu_gamma_beta0_pc
0.4 0.2 0.0 −0.2
0 2000 4000 6000 8000
log_sigma_gamma_beta0_pc
0.0 0.2 0.4 0.6 0.8 1.0
log_sigma_gamma_beta0_pc
1.00 0.75 0.50 0.25 0.00
0 2000 4000 6000 8000
mu_log_beta1
−2.4 −2.3 −2.2 −2.1 −2.0 −1.9 −1.8
mu_log_beta1
−1.8 −2.0 −2.2 −2.4
0 2000 4000 6000 8000
sigma_log_beta1
0.0 0.1 0.2 0.3 0.4 0.5 0.6
sigma_log_beta1
0.6 0.4 0.2 0.0
0 2000 4000 6000 8000
alpha_loglambda_p
alpha_loglambda_p
−2 −4

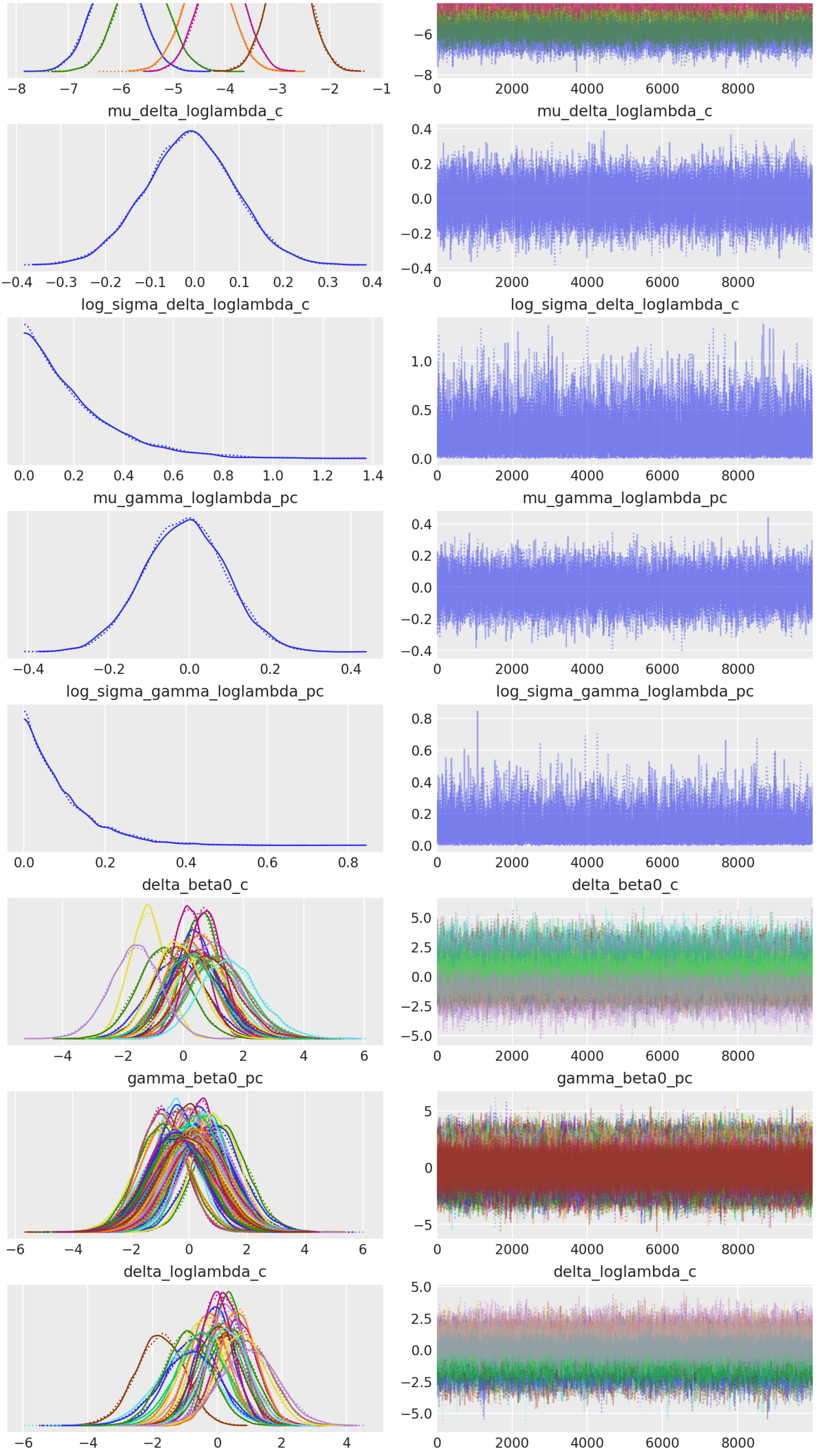
mu_delta_loglambda_c
mu_delta_loglambda_c
log_sigma_delta_loglambda_c
log_sigma_delta_loglambda_c
mu_gamma_loglambda_pc
mu_gamma_loglambda_pc
log_sigma_gamma_loglambda_pc
log_sigma_gamma_loglambda_pc
delta_beta0_c
delta_beta0_c
gamma_beta0_pc
gamma_beta0_pc
delta_loglambda_c
delta_loglambda_c

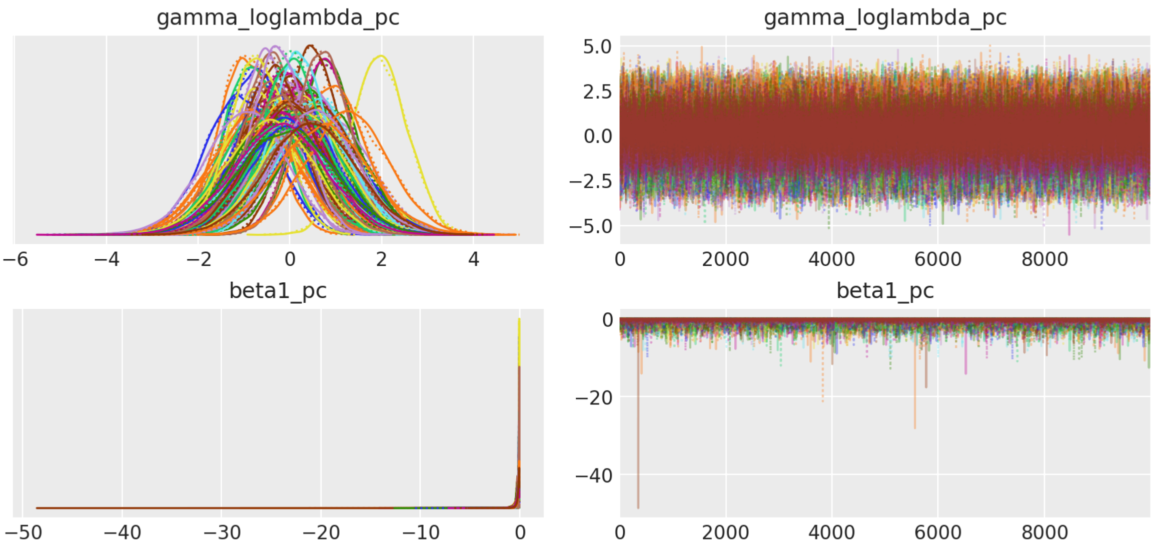


#### MODEL RESULTS

##### Results compilation

The results of the model primarily include the *posterior* distributions of the parameters, and the daily predation rate lambda, which accounts for the number of prey consumed per day per predator, and the two parameters underlying the digestion curve.

In [28]:
```
def Classeur_Estimations(path_to_save,
                         table_name,
                         df_dig,
                         df_pred,
                         path_to_InferenceData,
                         InferenceData,
                         N = 100) :

    Dataset = xr.open_dataset("".join([path_to_InferenceData, InferenceData]))
    Echantillons = az.convert_to_inference_data(Dataset)

    dict_values = {"Carabe":[], "Proie":[], "I_pc":[], "lambda_pc":[], "pI":[],
    lc = []; lp = []; lI = []; llamb = []; lpI = []; lfreq= []

    for cp, couple in enumerate(Echantillons.posterior.couples) :

        for i in range(N) :

            x = np.random.randint(0,len(Echantillons.posterior.draw))
            c = np.random.randint(0,1)

            Value_I_pc = np.asarray(Echantillons.posterior["I_pc"][c][x][cp])
            Value_lambda_pc = np.asarray(Echantillons.posterior["lambda_pc"][c][
            Value_pI = np.asarray(1 - np.maximum(0.001, np.exp(-(Value_lambda_pc

            dict_values["Carabe"].append(str(np.asarray(couple)).split("_")[0])
            dict_values["Proie"].append(str(np.asarray(couple)).split("_")[1])
            dict_values["I_pc"].append(Value_I_pc)
            dict_values["lambda_pc"].append(Value_lambda_pc)
            dict_values["pI"].append(Value_pI)
            dict_values["freq"].append(np.mean(df_pred[df_pred["Couple"] == coup

    df = pd.DataFrame(dict_values)
```

```
    df.to_excel("".join([path_to_save, table_name]))

    return df

try :

    df_Results = pd.read_excel("./Data/Estimates_IP_RFUII_0110.xlsx")

except :

    df_Results = Classeur_Estimations(path_to_save = "./Data/",
                                      table_name = "Estimates_IP_RFUII_0110.xlsx
                                      df_dig = df_dig,
                                      df_pred = df_pred,
                                      path_to_InferenceData = "./Samples/Echanti
                                      InferenceData = "Echantillon_Model_Digesti

liste_carabes = pd.unique(df_Results.Carabe)
df_Results = Arange_DescendingOrder(df_Results, df_pred)
```

Results display

***Digestion Curve***

The digestion curves for each predator-prey pair specified at the top of the cell (carabe = "PoCu" ; prey = "aphid") can be plotted in the cell below.

In [31]:
```
### Useful but uninteresting variables
Ech_t = np.arange(0,72,0.2)

liste_proies = [str(proie.values) for proie in list(Echantillons.posterior.proie
liste_carabes = [str(carabe.values) for carabe in list(Echantillons.posterior.ca
liste_couples = [str(couple.values) for couple in list(Echantillons.posterior.cc

dict_proies = {proie:i for i,proie in enumerate(liste_proies)}
dict_carabes = {carabe:i for i,carabe in enumerate(liste_carabes)}
dict_couples = {couple:i for i,couple in enumerate(liste_couples)}

dict_colors_prey = {"aphid" : "blue", "spider" : "orange", "springtail" : "red",
dict_colors_predator = {"PoCu" : "red", "BrSc" : "green", "AnDo" : "orange", "Am

def Figure_Digestion(dataset,
                     Echantillon,
                     proie,
                     carabe,
                     ax,
                     dict_color = dict_colors_predator) :

    df_Proie_par_Carabe = dataset.get_data(type_de_données = "labo",
                                           liste_carabes = [carabe],
                                           liste_proies = [proie],
                                           liste_variables = ["Carabe", "Tem

    couple = "_".join([carabe, proie])
    beta0_mean = np.mean(np.mean([np.asarray(Echantillon.posterior["alpha_beta0_
                       + np.mean([np.asarray(Echantillon.posterior["delta_beta0_c
                       + np.mean([np.asarray(Echantillon.posterior["gamma_beta0_p
    beta1_mean = np.mean([np.asarray(Echantillon.posterior["beta1_pc"])[1][i][di
```

```
    Δ_posterior_mean = [beta0_mean + beta1_mean*t for t in Ech_t]
    π_mean = [inv_logit(δ) for δ in Δ_posterior_mean]

    try :
        ax.plot(Ech_t, π_mean, color=dict_color[carabe], alpha=1, linewidth=2, l

    except :
        ax.plot(Ech_t, π_mean, color=dict_color[proie], alpha=1, linewidth=2, la

    for i in range(40) :

        r = np.random.randint(len(np.asarray(Echantillon.posterior["alpha_beta0_

        beta0_random = np.asarray(Echantillon.posterior["alpha_beta0_p".format(p
        beta1_random = np.asarray(Echantillon.posterior["beta1_pc"][1][r][dict_c

        Δ_posterior_random = [beta0_random + beta1_random*t for t in Ech_t]
        π_random = [inv_logit(δ) for δ in Δ_posterior_random]

        try :
            ax.plot(Ech_t, π_random, color=dict_color[carabe], alpha=0.1, linewi

        except :
            ax.plot(Ech_t, π_random, color=dict_color[proie], alpha=0.1, linewid
```

In [32]:

```
carabe = "PoCu" ; prey = "aphid"

fig = plt.figure(figsize=(5,5))
ax = fig.add_subplot(1,1,1)

Figure_Digestion(dataset,
                 Echantillons,
                 proie = prey,
                 carabe = "PoCu",
                 ax = ax,
                 dict_color = dict_colors_prey)

ax.tick_params(axis ='x',labelsize=10) ; ax.tick_params(axis ='y',labelsize=10)
ax.set_facecolor("white")
ax.set_xlabel("t : time after feeding (h)", fontsize=11)
ax.set_ylabel("$\pi (t)$ : probability of testing positive", fontsize=11)

plt.show()
```

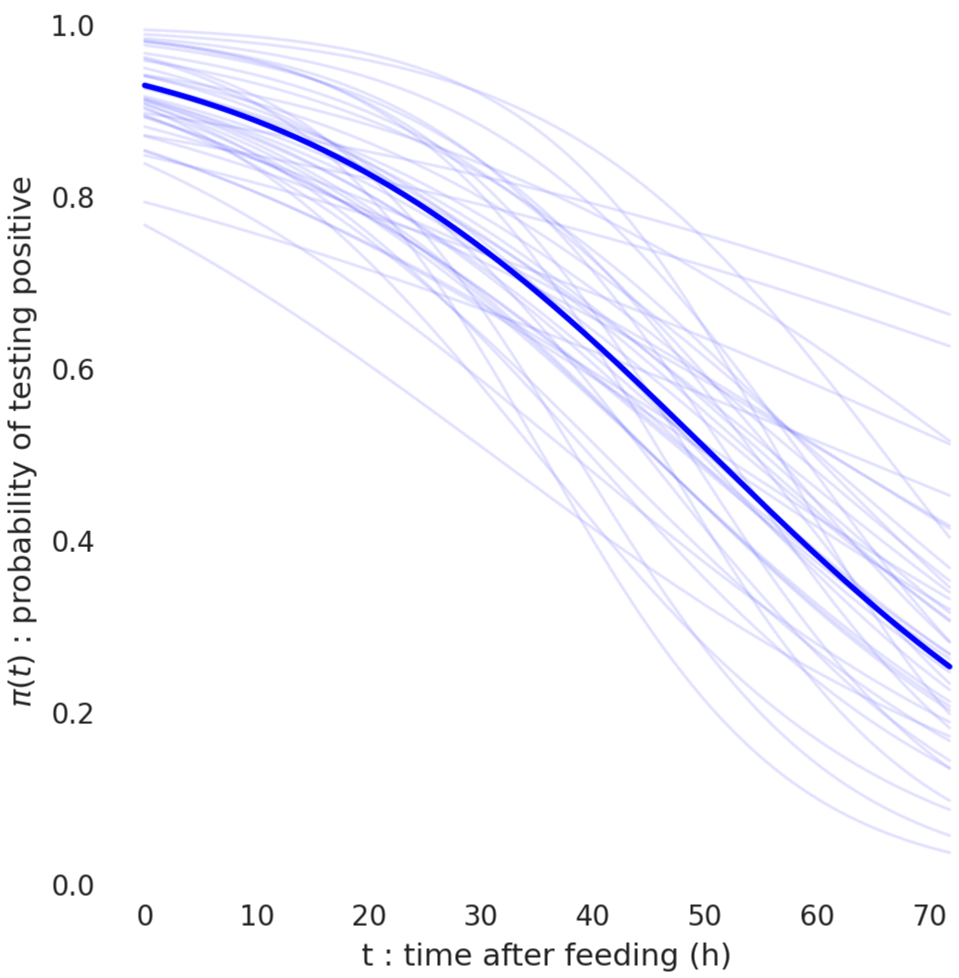


***Daily predation rate***

The cell below allows for plotting the daily predation rates posterior distributions for each predator-prey pair.

In [34]:
```
from Figures.Figures.Codes.Figures_Posteriors_Ridge_Plot import *

df_Predictions_sorted = Pre_traitement(df_Results,
                                       liste_carabes = pd.unique(df_Results.Cara
                                       param = "lambda_pc")


ridge_plot2(df_Results,
            list_predators = pd.unique(df_Results.Carabe),
            list_preys = pd.unique(df_Results.Proie),
            parameter = "lambda_pc")
```

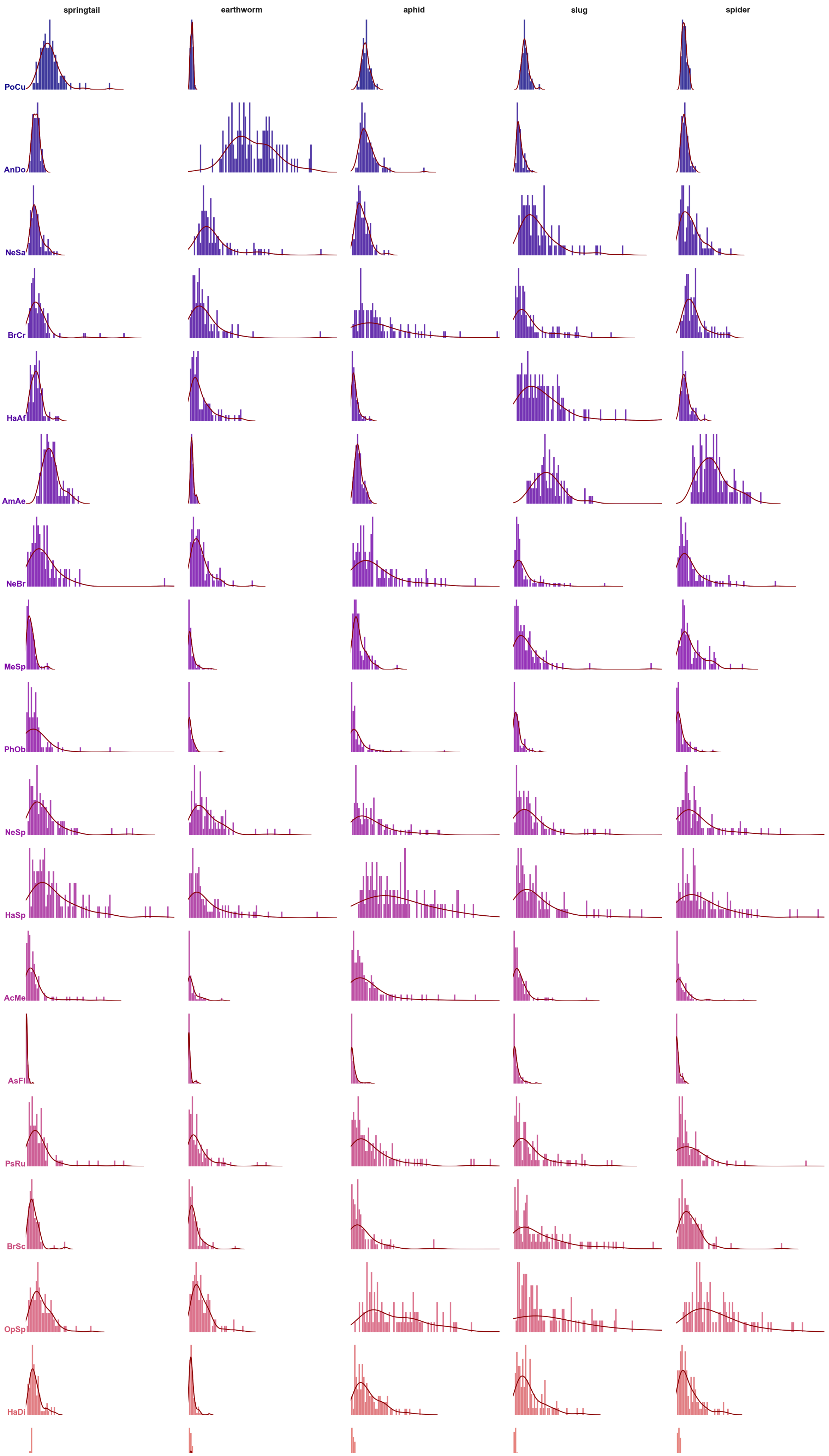
springtail
earthworm
aphid
slug
spider
PoCu
AnDo
NeSa
BrCr
HaAf
AmAe
NeBr
MeSp
PhOb
NeSp
HaSp
AcMe
AsFl
PsRu
BrSc
OpSp
HaDi

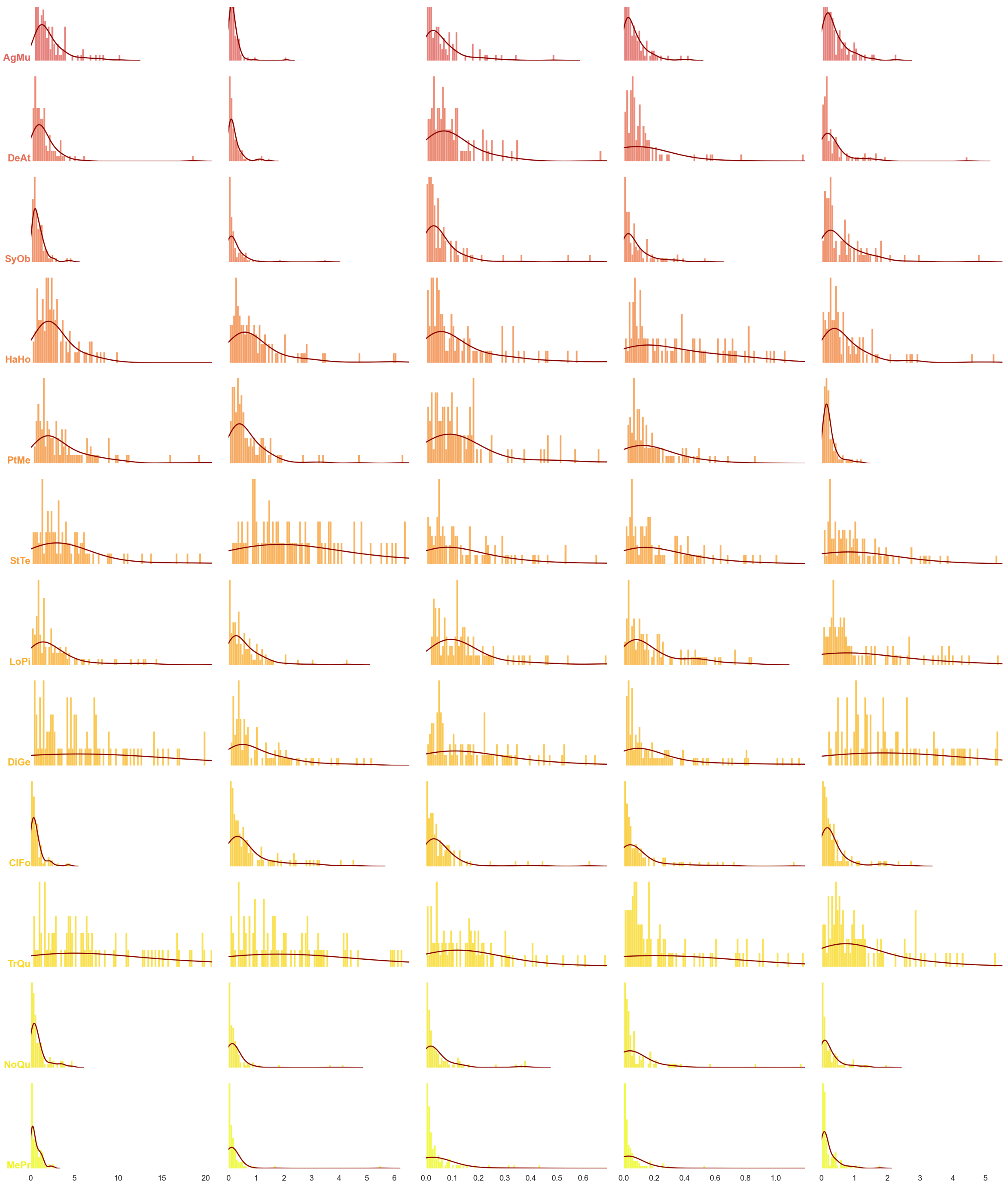


##### ***Daily predation rate***

The cell below allows for plotting the daily predation rates for every predator-prey pair.

In [36]:
```
fig, ax_ = plt.subplots(5, 1, figsize=(12,12))
dict_ylim = {"aphid" : 1.5, "earthworm" : 8, "slug" : 1.0, "spider" : 5, "spring

for i, proie in enumerate(["aphid", "earthworm", "slug", "spider", "springtail"]

    df_proie = df_Results[df_Results.Proie == proie]
    sns.boxplot(data = df_proie, x="Carabe", y="lambda_pc", ax = ax_[i],
                palette="dark", hue_order = pd.unique(df_proie.Carabe),
                showfliers=False, legend = False)

    ax_[i].set_ylim(-(dict_ylim[proie]/100), dict_ylim[proie])
    ax_[i].set_yticks([0, dict_ylim[proie]])
    ax_[i].set_yticklabels([0, dict_ylim[proie]], fontsize = 14)
```

```
    ax_[i].set_ylabel(proie, fontsize = 14)
    ax_[i].set_xlabel(None)
    ax_[4].set_xticklabels(pd.unique(df_proie.Carabe), fontsize = 10, rotation =

    if i != 4 :
        ax_[i].set(xticklabels=[])
        ax_[i].tick_params(bottom=False)

    ax_[4].set_xlabel("Carabid Species", fontsize = 14)
    if i != 0 :
        ax_[i].set_xlabel("")

plt.yticks(fontsize=14)
plt.show()
```

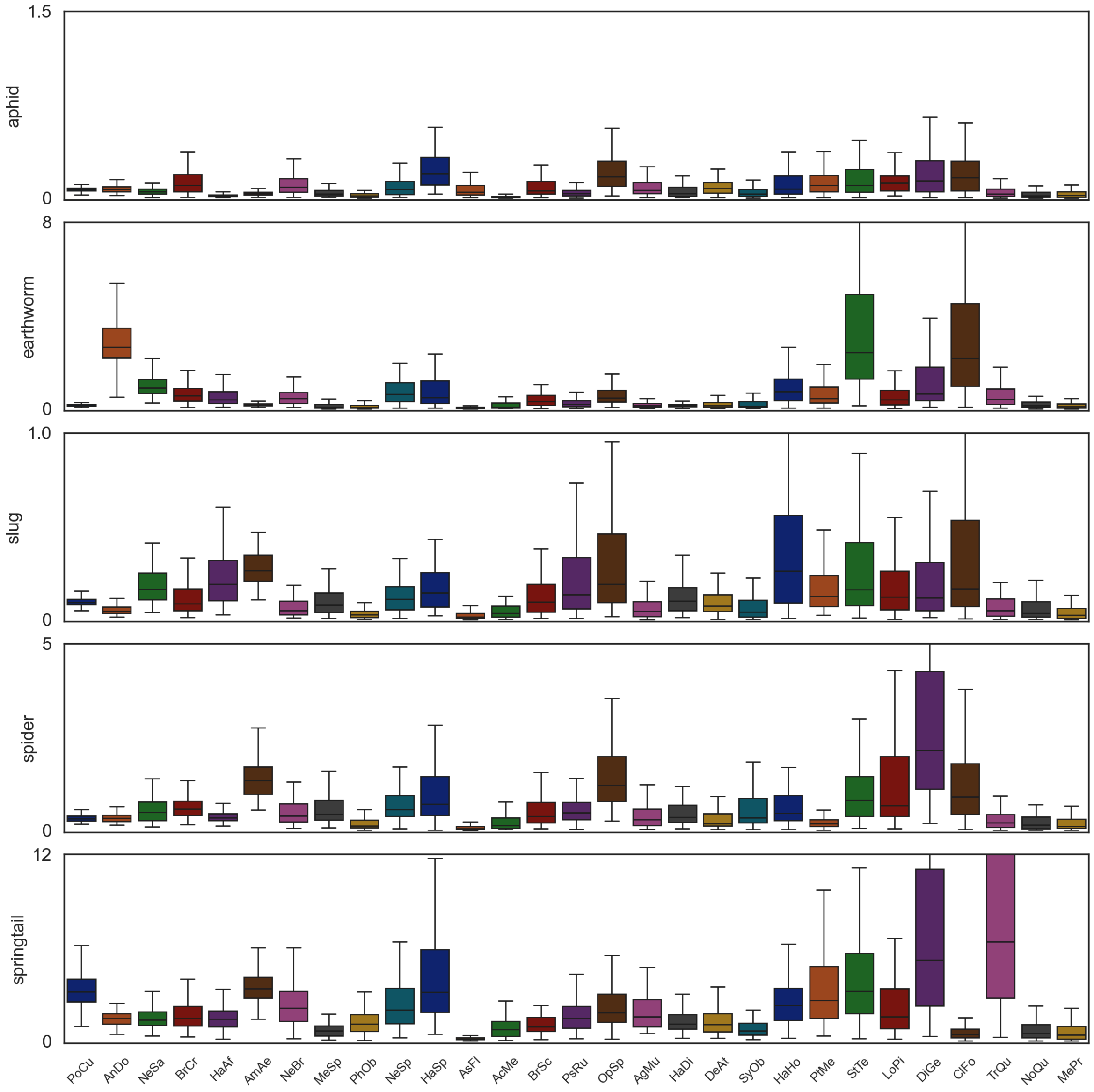


### Watermark

In [38]:
```
%load_ext watermark
%watermark -n -u -v -iv -w -p pytensor
```

```
Last updated: Tue Oct 01 2024

Python implementation: CPython
Python version       : 3.11.8
IPython version      : 8.22.2

pytensor: 2.12.3

seaborn   : 0.13.2
json      : 2.0.9
sklearn   : 1.4.1.post1
matplotlib: 3.8.3
platform  : 1.0.8
arviz     : 0.16.1
xarray    : 2024.2.0
scipy     : 1.12.0
re        : 2.2.1
pandas    : 2.2.1
sys       : 3.11.8 | packaged by conda-forge | (main, Feb 16 2024, 20:40:50) [MSC
v.1937 64 bit (AMD64)]
mycolorpy : 1.5.1
pymc      : 5.6.1
pytensor  : 2.12.3
numpy     : 1.25.2

Watermark: 2.4.3
```

---

■ *End of document*
